# Liquid-like transcription condensates locally constrain chromatin in living human cells

**DOI:** 10.1101/2025.07.05.663270

**Authors:** Adilgazy Semeigazin, Katsuhiko Minami, Masa A. Shimazoe, Saadi Khochbin, Daniel Panne, Satoru Ide, Kazuhiro Maeshima

**Author notes:** Correspondence, National Institute of Genetics, Mishima, Shizuoka 411-8540, Japan. Cell Biology Center, Institute of Integrated Research, Institute of Science Tokyo (Satoru Ide).

## Abstract

The organization and dynamics of chromatin play important roles in transcriptional regulation. The transcription machinery is known to constrain chromatin dynamics. Recently, transcription condensates formed via liquid–liquid phase separation (LLPS) or other mechanisms have emerged as key regulators of gene expression. What is the physical nature of such condensates in the cell? Do they interact with and constrain chromatin? To address these questions, we focused on BRD4-NUT, a fusion oncoprotein found in NUT carcinoma. Using single-molecule dual-color imaging, we found that individual BRD4-NUT molecules diffuse within condensates like a viscous liquid in live human cells. Single-nucleosome imaging specific to euchromatin shows that these liquid-like condensates restrict the movement of chromatin through BRD4 bromodomain-dependent crosslinking of neighboring acetylated nucleosomes. Our findings uncover a previously unrecognized mechanism by which LLPS-based condensates modulate chromatin dynamics, suggesting that condensates contribute to genome regulation through physical, and not only biochemical, control.

## Introduction

Genomic DNA is wrapped around core histones (two copies each of H2A, H2B, H3, and H4) to form nucleosomes ^1–3^. In higher eukaryotic cells, a string of nucleosomes, together with other proteins and RNAs, is organized into chromatin domains ^4,5^. Within these domains, nucleosomes fluctuate like a liquid ^6,7^, facilitating genome functions. Transcriptional regulation in eukaryotic cells occurs within this complex, dynamic, and crowded chromatin environment ^8–11^.

Recent studies have demonstrated that transcription factors and coactivators can form biomolecular condensates ^11–17^. In particular, bromodomain-containing protein 4 (BRD4), Mediator, and RNA polymerase II (RNA Pol II) form such condensates at super-enhancers, which play regulatory roles in gene expression. These condensates are proposed to form via liquid–liquid phase separation (LLPS) ^12,13,18^, which may facilitate efficient transcription by concentrating active components and excluding inhibitory factors ^19,20^.

While LLPS-driven transcriptional condensates have attracted broad interest, concrete evidence supporting this concept remains limited, especially in living cells ^21,22^. Although RNA Pol II, BRD4, and several transcription factors form droplets or condensates *in vitro*, it remains unclear whether similar structures truly arise in cells through LLPS or via alternative mechanisms. A major challenge lies in the lack of robust methods to test this concept in live-cells.

For example, treatment with 1,6-hexanediol (1,6-HD), which can dissolve LLPS-driven droplets *in vitro* and *in vivo*, is a frequently used approach ^21^. However, 1,6-HD treatment also rapidly immobilizes and condenses chromatin in living cells—a distinct effect that is unrelated to its reported droplet-disrupting activity—suggesting that interpretations of its effects should be made with caution ^23^. Fluorescence recovery after photobleaching (FRAP) is widely used to assess the mobility of fluorescently tagged proteins within droplets or condensates. However, FRAP alone does not directly determine whether these structures exhibit liquid-like properties ^21,24^. Higher-resolution quantitative analyses of liquid droplets/condensates in living cells are needed to examine whether their component molecules diffuse like a liquid or not. To this end, single-molecule imaging and tracking is a promising approach ^21^.

Beyond their formation mechanisms, how condensates in cells interact with and impact the underlying chromatin structure and dynamics remains largely unclear. There is considerable evidence indicating that RNA Pol II depletion or inhibition enhances chromatin movement ^25–28^ ^29,30^. While this suggests that transcription machinery somehow constrains chromatin, it is still poorly understood whether transcriptional condensates themselves influence the physical behavior of chromatin in their vicinity for transcriptional regulation.

To approach these two issues, BRD4-NUT, a fusion oncoprotein found in NUT carcinoma (NC), provides a unique model system ^31,32^. BRD4-NUT forms prominent nuclear condensates via the tandem bromodomains of BRD4 and the p300-interacting and activating NUT region, leading to the formation of hyperacetylated chromatin foci ^33–36^ or Hi-C “megadomains”^37^.

In this study, we developed single-molecule dual-color imaging ^6,7^ to investigate the biophysical and functional properties of BRD4-NUT condensates in live cells. We demonstrate that individual BRD4-NUT molecules fluctuate within condensates in a liquid-like manner.

Combined with euchromatin-specific single-nucleosome imaging ^38^, which can sensitively detect changes in chromatin state, we reveal that these liquid-like condensates physically constrain the motion of nearby nucleosomes in active chromatin regions. This effect was observed both in engineered HeLa cells expressing BRD4-NUT and in NC-derived human HCC2429 cells. Furthermore, we demonstrate that the nucleosome-constraining activity is mediated by the BRD4S (short isoform of BRD4) moiety of BRD4-NUT via tandem bromodomain-dependent interactions with acetylated nucleosomes. Together, our results uncover a previously unappreciated role of BRD4-driven condensates in modulating local chromatin dynamics and suggest a general mechanism by which LLPS-based transcriptional regulators constrain chromatin mobility to regulate gene expression.

## Results

### BRD4-NUT condensate formation upon doxycycline induction in HeLa cells

We focused on BRD4-NUT as a model transcription condensate ^35,36^ to explore the interplay between condensates and chromatin. In NC cells, the BRD4-NUT protein arises from an in-frame fusion between the coding regions of the *BRD4* and *NUTM1* genes ^31,33^ and is not normally expressed in non-NC cells, such as HeLa. To avoid potential cytotoxicity of BRD4-NUT in HeLa cells, we generated a doxycycline-inducible EGFP-tagged BRD4-NUT expression system using Tet-On 3G technology (Figs. 1A(i) and S1; also see Fig. S3) ^39,40^. This system was stably integrated into the AAVS1 locus on chromosome 19—considered a genomic safe harbor ^41^—by CRISPR/Cas9-mediated genome editing (Fig. S1).

**Fig. 1.**
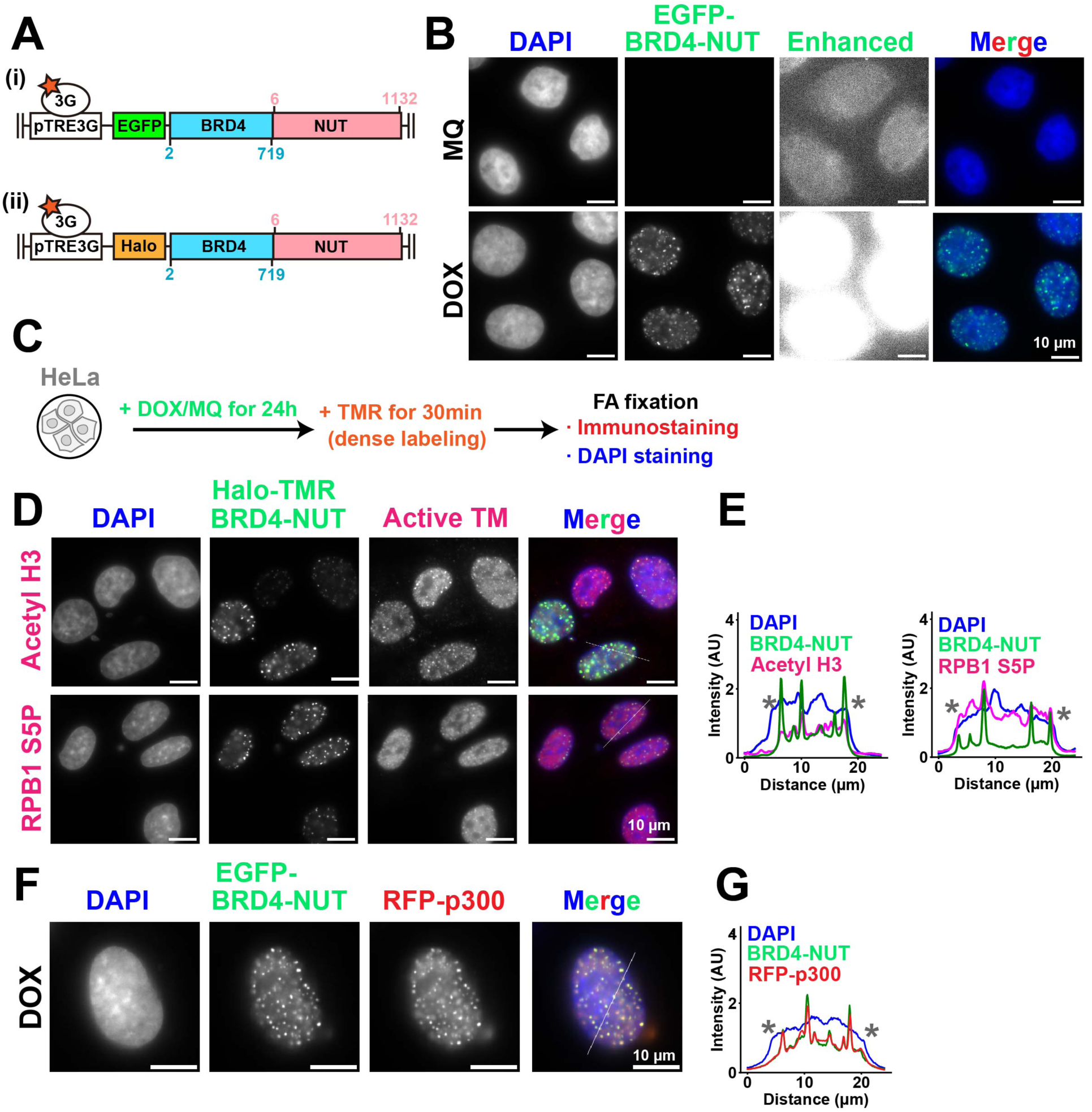
BRD4-NUT forms condensates that colocalize with transcription proteins. **(A)** Schematic of the Tet-On 3G for doxycycline-inducible ectopic expression of (i) EGFP- or (ii) Halo-BRD4-NUT. **(B)** HeLa cells treated either with 0.1% Milli-Q (MQ) water or 1 µg/mL doxycycline (DOX) for 24 h to express EGFP-BRD4-NUT before FA fixation. Note that EGFP-BRD4-NUT foci are only seen with DOX but not with MQ, even when significantly enhanced (Enhanced). Scale bars, 10 μm. The same intensity scale was used for all images in each of the columns. **(C)** Simplified protocol for the observation of BRD4-NUT condensates in FA-fixed cells. **(D)** Immunostaining against pan-acetylated histone H3 (Acetyl H3; upper row) or RNA Pol II phosphorylated at Ser-5 (RPB1 S5P; lower row) in HeLa cells expressing Halo-BRD4-NUT with DOX treatment. Scale bars, 10 μm. Image intensities were optimized for visualization. **(E)** Line plot as derived from the thin white line crossing the nucleus from (D). Blue, DAPI (DNA); magenta, Active Transcription Machinery (Active TM for Acetyl H3 (left) or RPB1 S5P (right)); green, Halo-TMR-BRD4-NUT. Note a significant overlap between Halo-TMR-BRD4-NUT or Acetyl H3 and RPB1 S5P signal peaks. Asterisks (**\***) show the nuclear edge positions. **(F)** HeLa cells transiently co-transfected with EGFP-BRD4-NUT and RFP-p300 plasmids and treated with 1 µg/mL doxycycline for 24 h before FA fixation. Note that RFP-p300 forms condensates in the presence of EGFP-BRD4-NUT. Scale bars, 10 μm. **(G)** Line plot as derived from the thin white line crossing the nucleus from (F). Blue, DAPI (DNA); red, RFP-p300; green, EGFP-BRD4-NUT. Note a significant overlap between EGFP-BRD4-NUT and RFP-p300 signal peaks. Asterisks (**\***) show the nuclear edge positions.

To assess the localization of EGFP-BRD4-NUT in HeLa cells, the cells were subjected to a 24-hour induction with doxycycline (see Materials and Methods for details). The uninduced control cells did not show any GFP signal, which confirmed there was not a significant leakage of EGFP-BRD4-NUT expression (upper, Fig. 1B). Most cells exposed to doxycycline exhibited bright, discrete EGFP-BRD4-NUT nuclear foci (lower, Fig. 1B; also Fig. 1F), closely resembling the BRD4-NUT condensates reported previously ^33,35,36^. The expression levels of EGFP-BRD4-NUT, as well as the morphology of the condensates, were highly variable among cells. The EGFP-BRD4-NUT condensates often displayed markedly different brightness, sizes, and shapes. Using a similar system (Fig. S1), we expressed HaloTag-fused BRD4-NUT (Halo-BRD4-NUT) (Fig. 1A(ii); also see Fig. S3). When we induced expression for 24 hours and labeled Halo with an excess amount (5 nM) of tetramethylrhodamine (TMR; Fig. 1C), we observed large nuclear foci of Halo-TMR-BRD4-NUT (Fig. 1D), comparable to those of EGFP-BRD4-NUT (Fig. 1B). To determine whether active transcription machineries formed co-condensates with Halo-BRD4-NUT, pan-acetylated histone H3 (H3Ac; upper, Fig. 1D) and Ser5-phosphorylated RNA Pol II (RPB1-S5P, the largest subunit of RNA Pol II) (lower, Fig. 1D) were detected by immunostaining. The localization of p300 was assessed by transient ectopic co-expression of RFP-tagged p300 (Fig. 1F) and EGFP-BRD4-NUT. Both H3Ac (upper, Fig. 1D; left, Fig. 1E) and p300 (Fig. 1F-G) showed clear signal enrichment within BRD4-NUT foci, consistent with previous reports ^33,42^. RPB1-S5P, an active RNA Pol II marker, also had signal peaks overlapping with BRD4-NUT (right, Fig. 1E).

Since BRD4-NUT recognizes acetylated nucleosomes, we examined whether BRD4-NUT condensates are dispersed by using the histone deacetylase (HDAC) inhibitor trichostatin A (TSA), which increases genome-wide histone acetylation levels ^43^. EGFP-BRD4-NUT was expressed for 24 hours and the cells were treated with TSA in the last 8 hours (Fig. S2A). The TSA treatment almost completely dispersed BRD4-NUT condensates across the nucleus (Fig. S2B, C). These results indicate that the induced Halo-BRD4-NUT condensates have properties closely matching those reported previously ^33,35,36^.

### Single molecules of BRD4-NUT fluctuate inside the condensates

While a recent fluorescence recovery after photobleaching (FRAP) experiment pointed to a rapid internal turnover of BRD4-NUT condensates in cells ^35^, whether BRD4-NUT molecules diffuse like a liquid within these condensates remains unclear. To approach this issue, we performed single-molecule BRD4-NUT imaging inside these condensates. To observe both single molecules and whole condensates of Halo-BRD4-NUT in the same cell (left, Fig. 2A), we simultaneously labeled Halo with a low concentration of JF646 (50 pM) and a high concentration of TMR (5 nM) (Fig. 2B). Using oblique illumination microscopy (right, Fig. 2A) ^44–46^, sparse and dense labeling of the condensates allowed individual Halo-BRD4-NUT dots and whole condensates to be clearly observed, respectively (Fig. 2C(i and iii); Movie S1). We acquired continuous image sequences of the dots and condensates at 10 ms per frame (500 frames, 5.0 s total).

**Fig. 2.**
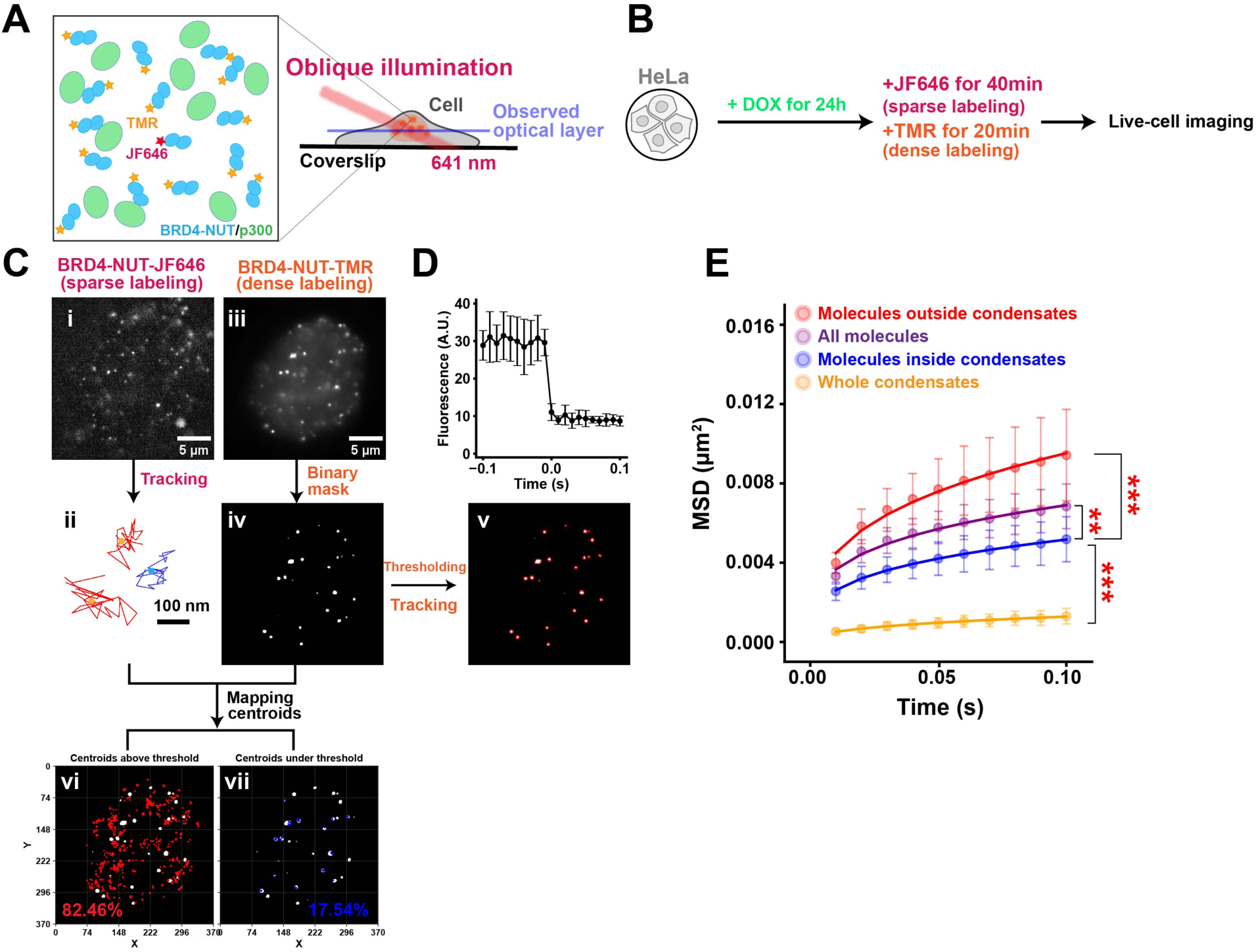
Single molecules of BRD4-NUT are trapped by and fluctuate relative to the condensates. **(A)** Left: cartoon representation of a cell expressing Halo-BRD4-NUT/p300 condensates in the nucleus. Note that HaloTag can be simultaneously labeled with different fluorophores (e.g. JF646 and TMR). Right: oblique illumination microscopy. The laser (red) can excite fluorescent molecules within a limited thin optical layer (blue) of the nucleus and reduce background noise. Only the 641-nm laser is indicated. **(B)** Simplified protocol for sparsely labeling molecules inside Halo-BRD4-NUT condensates in live HeLa cells. **(C)** Workflow of the analysis of single-molecule motions “inside” and “outside” BRD4-NUT condensates. i: Image of Halo-BRD4-NUT sparsely labeled with JF646 after background subtraction. ii: Representative trajectories of three tracked single molecules. Blue, “inside”; Red, “outside”. Trajectory centroids are indicated with circles. iii: Image of Halo-BRD4-NUT densely labeled with TMR in the same cell as i. iv: Representative binary mask obtained from Halo-TMR-BRD4-NUT image in iii. v: Example for the detection of condensates from the binary mask in iv for tracking. vi: centroids of “outside” Halo-JF646-BRD4-NUT trajectories overlayed on the binary mask from iv. % - their percentage from all trajectories. vii: centroids of “inside” Halo-JF646-BRD4-NUT trajectories overlayed on the binary mask from iv. % - their percentage from all trajectories. **(D)** Single-step photobleaching profile of ten Halo-JF646-BRD4-NUT dots. Mean ± SD among 10 dots is shown. The vertical axis represents fluorescence intensity. **(E)** MSD plots (± SD among cells) of Halo-BRD4-NUT all (purple), “inside” (blue), and “outside” (red) JF646 single molecules, and TMR whole condensates (orange) in a short time range (0.01 to 0.1 s). For each sample, *n* = 20 cells. ***, *p* = 1.1 × 10^−8^ for “inside” versus “outside”; *p* = 1.5 × 10^-11^ for “inside” versus “whole condensates”; **, *p* = 4.0 × 10^-3^ for “inside” versus “all” single molecules by two-sided Kolmogorov-Smirnov test. Note that MSD of “inside” molecules is significantly smaller than that of both “outside” and “all”, but larger than “whole condensates”.

Each fluorescent dot of JF646 displayed a single-step photobleaching pattern (Fig. 2D), a characteristic of single molecules. We first detected Halo-BRD4-NUT dots in each image using the LoG detector of TrackMate (Fig. 2C(ii)) ^47^ and fitted them to an assumed Gaussian function. The trajectories of each dot were then tracked for 500 frames (5 s total). The position determination accuracy of Halo-BRD4-NUT was 16.5 nm (Fig. S4A). We focused on the movements of molecules primarily associated with the condensates under these imaging and tracking conditions, while omitting very fast trajectories. To track whole condensates, each frame from 1 to 100 of the condensate movies was binarized (Fig. 2C(iv)), and the condensate boundaries were detected using the Thresholding detector of TrackMate (Fig. 2C(v)). The trajectory centroids of Halo-JF646-BRD4-NUT were then mapped onto the binary masks of the Halo-TMR-BRD4-NUT condensates and categorized as “inside” or “outside” the condensates (Fig. 2C(vii) or (vi) respectively).

Observed Halo-JF646-BRD4-NUT single-molecule motion inside the condensates exhibited about 60 nm of displacement over 20 ms of observation time (Fig. S4B). Their calculated mean squared displacement (MSD) showed subdiffusive motion of Halo-JF646-BRD4-NUT (blue, Fig. 2E), which was significantly lower than that of single molecules outside the condensates (red, Fig. 2E), suggesting that the condensates confine the movement of BRD4-NUT molecules trapped inside them. Interestingly, the motion of Halo-JF646-BRD4-NUT molecules inside the condensates (blue, Fig. 2E) was much lower than the averaged nucleosome motion reported previously ^48^. Nevertheless, the motion of Halo-JF646-BRD4-NUT single molecules inside the condensates was much more dynamic than that of whole condensates (Halo-TMR-BRD4-NUT) (orange, Fig. 2E). The diffusion coefficient of single molecules (*D* = 1.7 × 10^-3^ µm²/s) was greater than that of whole condensates (*D* = 5.3 × 10^-4^ µm²/s) in the time range between 0 and 0.1 s (Fig. S4C). Furthermore, the radius of confinement (*R*_c_) ^49^ of single molecules inside the condensates and whole condensates were estimated to be 84.02 ± 14.58 nm and 55.38 ± 14.26 nm, respectively (Fig. S5A). Similar trends were observed at an exposure time of 50 ms/frame (Fig. S5B; Movie S2) for more extended tracking periods. Taken together, all our findings suggest that the single molecules of BRD4-NUT fluctuate inside the condensates in which they reside.

### BRD4-NUT condensates exhibit viscous liquid-like behavior

Simple single-nucleosome imaging does not reveal how much individual BRD4-NUT molecules fluctuate within the condensate because the condensate itself moves (orange, Fig. 2E; orange, Fig. S5B). To overcome this problem, we focused on the distances between two neighboring BRD4-NUT molecules in the condensate (<300 nm distance, Figs. 3A-B and S6A) and analyzed the mean squared distances between them (two-point MSD) ^6,7,50^ because the distance between molecules is much less affected by either translational or rotational movements of the condensate during imaging.

**Fig. 3.**
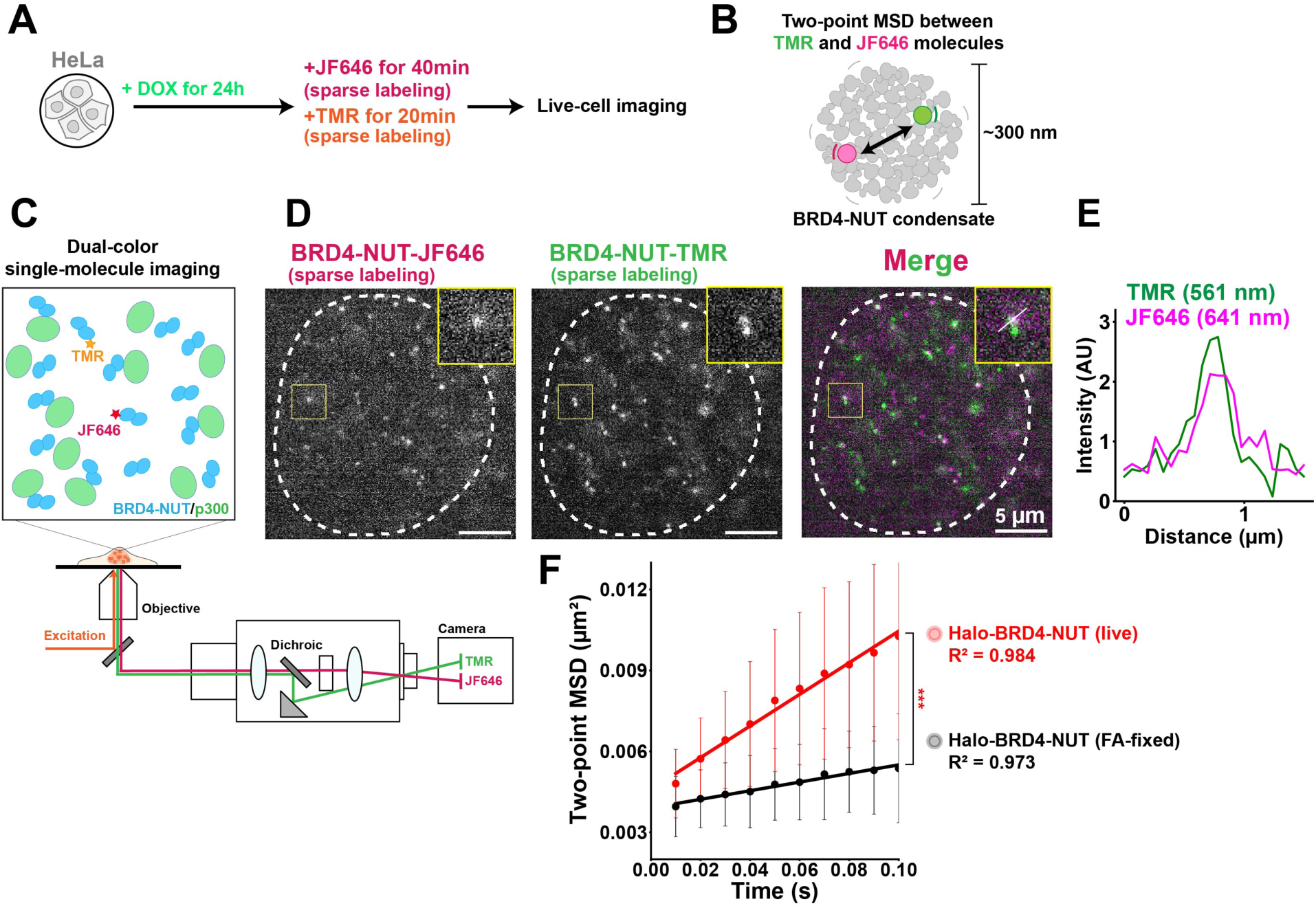
BRD4-NUT condensates exhibit viscous liquid-like physical properties. **(A)** Simplified protocol for sparsely labeling Halo-BRD4-NUT molecules with ligands (JF646 and TMR) inside their condensates in live HeLa cells. **(B)** Schematic for two-point MSD analysis. The mean square relative distances between the TMR and JF646 molecules were tracked. The neighboring dot pairs whose averaged distances were <300 nm were analyzed. **(C)** Schematic for dual-color imaging with a beam splitter system (W-VIEW GEMINI, Hamamatsu Photonics). The images of two single molecules with different colors were simultaneously acquired with a single sCMOS. Note that while many molecules were left unlabeled, fluorescent molecules were only singly labeled with either JF646 or TMR. **(D)** Representative images of neighboring single molecules labeled with JF646 (left) and TMR (center) in living HeLa cells. Right: the merged image. Representative single molecules are selected in yellow boxes and shown separately (top right corner). A white punctate line delineates the nucleus. The intensity line profiles for TMR and JF646 across the white line in the last yellow box in (D) and is plotted in **(E)**. **(F)** Two-point MSD plots (mean ± SD among cells) between Halo-JF646- and Halo-TMR-BRD4-NUT single molecules in live (red; *n* = 28 cells) and fixed (black; *n* = 28 cells) HeLa cells from 0.01 to 0.1 s (tracked at 10 ms/frame,). ***, *p* = 3.7 × 10^−7^ for “live” versus “fixed” by two-sided Kolmogorov-Smirnov test.

For this purpose, we developed dual-color single-molecule imaging to evaluate the two-point MSD between pairs of spatially neighboring BRD4-NUT molecules labeled with TMR and JF646 (Fig. 3A and C). Using a beam splitter system to acquire single-molecule dual color movies at 10 ms/frame for 500 frames (Fig. 3C) ^6,7^, we simultaneously observed clear dots in each color channel as single molecules (Figs. 3D, S6B-F; Movie S3). The position determination accuracy was 16.3 nm for TMR and 14.3 nm for JF646 (Fig. S6C and E). Notably, there was essentially no cross-talk between the TMR and JF646 signals (Figs. 3D-E, S6F-G). Among these, we found many pairs of Halo-BRD4-NUT molecules with TMR and JF646 signals located in close proximity (Fig. 3D; upper, Fig. S6F; upper, Fig. S6G).

We collected Halo-BRD4-NUT pairs whose distance was less than 300 nm and analyzed them. This distance threshold was chosen because live-cell condensate images, on average, showed an estimated diameter of ∼6 pixels (∼390 nm; Fig. S6A), suggesting that such pairs likely reside within the same condensates. Strikingly, the two-point MSD plot showed a nearly perfect linear relationship (Fig. 3F, *R*² = 0.984), suggesting that single molecules of Halo-BRD4-NUT diffuse inside the condensates like a viscous liquid. Importantly, when we tracked the single molecules at 50 ms/frame in the time range between 0.05 and 0.5 s, the two-point MSD plot was again linear (Fig. S6H, *R*² = 0.985; Movie S4) while the two-point MSD plot of two neighboring nucleosomes (<150 nm apart) exhibited subdiffusive behavior ^6^. These findings provide direct evidence that BRD4-NUT condensates form via LLPS and possess viscous liquid-like physical properties.

### BRD4-NUT condensates locally constrain the motion of euchromatic nucleosomes

How do liquid-like BRD4-NUT condensates interplay with chromatin? To address this question, we performed euchromatin-specific single-nucleosome imaging after expressing EGFP-BRD4-NUT for 24 hours, and analyzed the nucleosome trajectories around the condensates (Fig. 4A, B). Since ectopically expressed H4-Halo is incorporated into nucleosomes mainly in euchromatic regions with active chromatin marks (Hi-C A-compartment) (Fig. S7, ^38^), we took advantage of this for euchromatin-specific single-nucleosome imaging.

**Fig. 4.**
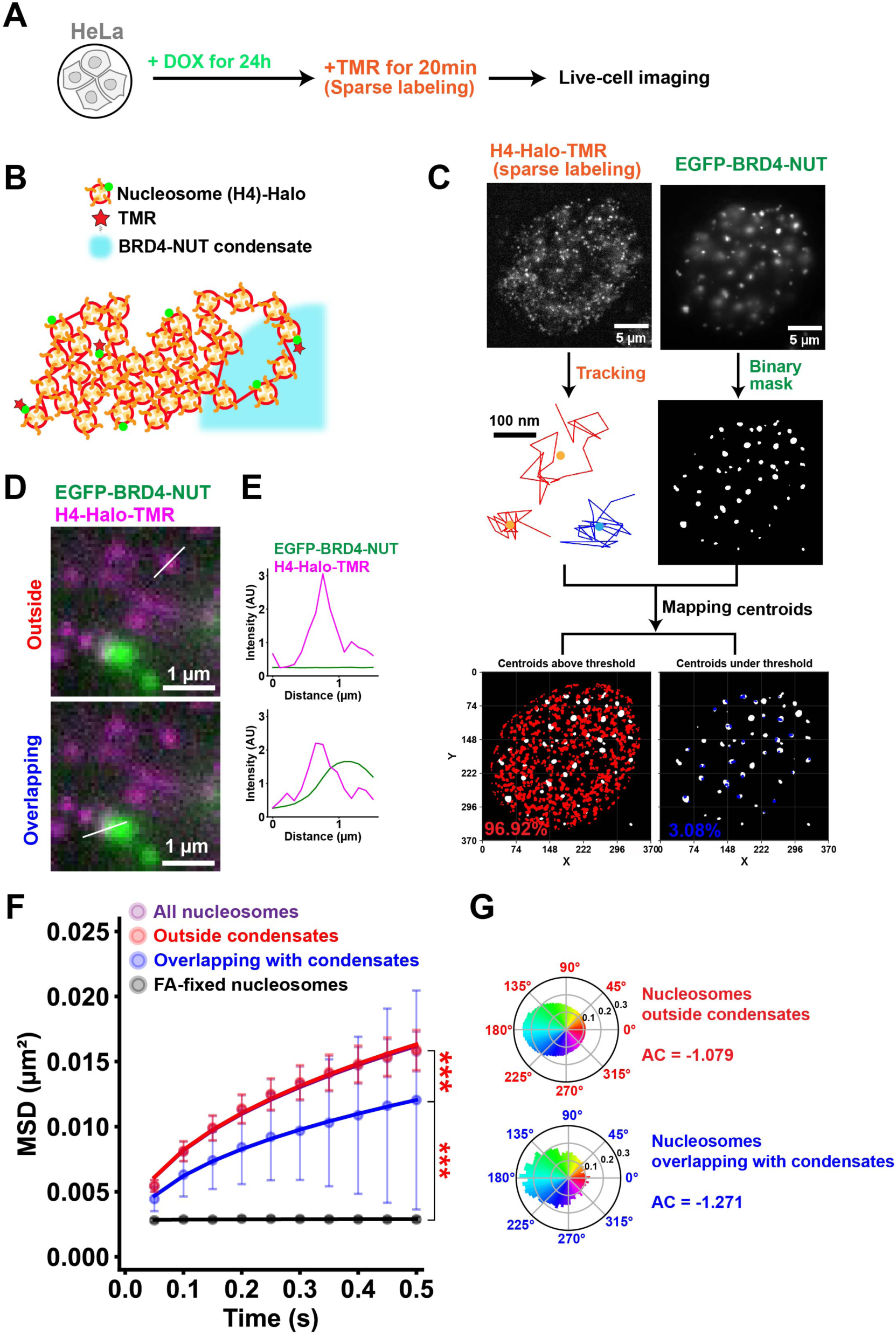
BRD4-NUT condensates locally constrain the motion of euchromatic nucleosomes. **(A)** Simplified protocol for sparsely labeling H4-Halo nucleosomes in live HeLa cells expressing EGFP-BRD4-NUT. **(B)** A small fraction of H4-Halo was fluorescently labeled with TMR (red star) for imaging single nucleosomes overlapping or not overlapping with BRD4-NUT condensates (cyan sphere). **(C)** Workflow of the analysis of single-nucleosome motions overlapping and not overlapping (“outside”) with EGFP-BRD4-NUT condensates. Top left: sparsely labeled H4-Halo image after background subtraction. Middle left: representative trajectories of three tracked single nucleosomes. Blue, “overlapping”; Red, “outside”. Trajectory centroids are indicated with circles. Top right: EGFP-BRD4-NUT image in the same cell as top left. Middle right: representative binary mask obtained from EGFP-BRD4-NUT image in top right. Bottom left: centroids of “outside” H4-Halo trajectories overlayed on the binary mask from middle right. % - their percentage from all trajectories. Bottom right: centroids of “overlapping” H4-Halo trajectories overlayed on the binary mask from middle right. **(D)** Example overlays of H4-Halo-TMR nucleosomes (magenta) outside (upper row) or overlapping with (lower row) with EGFP-BRD4-NUT condensates (green). The intensity line profiles for TMR and EGFP across the white lines in (D) are plotted in **(E)**. **(F)** MSD plots (± SD among cells) of H4-Halo nucleosomes in live (purple) and FA-fixed (black) cells. MSD plots (± SD among cells) of “overlapping” (blue) and “outside” (red) H4-Halo nucleosomes in live cells. For each sample, *n* = 25 cells. ***, *p* = 2.5 × 10^−7^ for “overlapping” versus both “outside” and “all” H4-Halo trajectories; *p* = 1.6 × 10^−14^ for “overlapping” versus FA-fixed nucleosomes by two-sided Kolmogorov-Smirnov test. **(G)** Measured angle distribution of “overlapping” and “outside” nucleosomes (607,929 total number of angles for “outside”; 14,758 angles for “overlapping”).

H4-Halo was sparsely labeled with TMR (top left, Fig. 4C) and observed by oblique illumination microscopy (right, Fig. 2A) ^44–46^. The clearly resolved individual H4-Halo-TMR dots were imaged at 50 ms/frame (top left,Fig. 4C; left, Movie S5) and tracked. At this frame rate, we could focus on the behavior of nucleosomes labeled by H4-Halo-TMR ^38^. Their trajectories were extracted as a set of (*x, y*) coordinates of their Gaussian centroids (middle left, Fig. 4C). The EGFP-BRD4-NUT condensates were separately captured in a different channel (top right, Fig. 4C). The EGFP-BRD4-NUT images were binarized as described previously (middle right, Fig. 4C) ^6^. In the resulting masks, the EGFP-BRD4-NUT condensates appeared white, while the background was black. The centroids of H4-Halo trajectories were mapped onto the binary masks and categorized based on whether they belonged to the white or black regions (bottom, Fig. 4C) ^51^. If a centroid localized to a white region of the mask, the nucleosome trajectory was considered to overlap with the condensates (lower, Fig. 4D-E). This group of nucleosomes is usually a minority (<5%) and positioned adjacent to the condensates, which may affect their dynamic behavior. Alternatively, if a centroid localized to a black region, the trajectory was considered to reside outside the condensates (upper, Fig. 4D-E). This group includes the majority of nucleosomes (>95%).

When the trajectory grouping was complete, the nucleosome MSDs (Fig. 4F) and asymmetry coefficients (AC) ^52^ (Fig. 4G) of motion angle distributions (motion vectors) were calculated and compared. The AC value indicates how much pulling-back force the nucleosomes experience; that is, a smaller AC value shows more constraint (Fig. S8) ^48^. Strikingly, the nucleosome motion whose trajectories overlapped with the EGFP-BRD4-NUT condensates was more constrained than that outside of them (Fig. 4F, G) and became closer to that of BRD4-NUT molecules inside the condensates (blue, Fig. S5B), suggesting the condensates can restrict the thermal fluctuation of nearby nucleosomes. To validate the result, we assigned the H4-Halo trajectory centroids to randomized masks (Fig. S9A) and re-calculated the MSDs and ACs (Fig. S9B-C). In this case, there was no difference between nucleosome motions overlapping with the wrong masks and those outside (Fig. S9B-C), excluding the possibility that the result is an artifact of the categorization. Taken together, we conclude that the EGFP-BRD4-NUT condensates indeed constrain the motion of neighboring nucleosomes.

We then focused on BRD4-NUT in NC cells ^33^. Because endogenous BRD4-NUT does not have any fluorescent tag, we first marked the condensates in an NC cell line, HCC2429, by constitutive ectopic expression of EGFP-BRD4-NUT (Fig. 5A-D). The intensity of BRD4-NUT foci was much lower in HCC2429 cells than in HeLa cells (Fig. S10A), which likely reflects the endogenous BRD4-NUT condensates. Consistently, immunostaining revealed that the intensity of endogenous BRD4-NUT condensates in HCC2429 parental cells was approximately 6.7 times lower than that of EGFP-BRD4-NUT condensates in HeLa cells (Fig. S10B), and was comparable to that of BRD4-NUT condensates in HCC2429 cells expressing EGFP-BRD4-NUT (only ∼1.3 times difference; Fig. S10C, D).

**Fig. 5.**
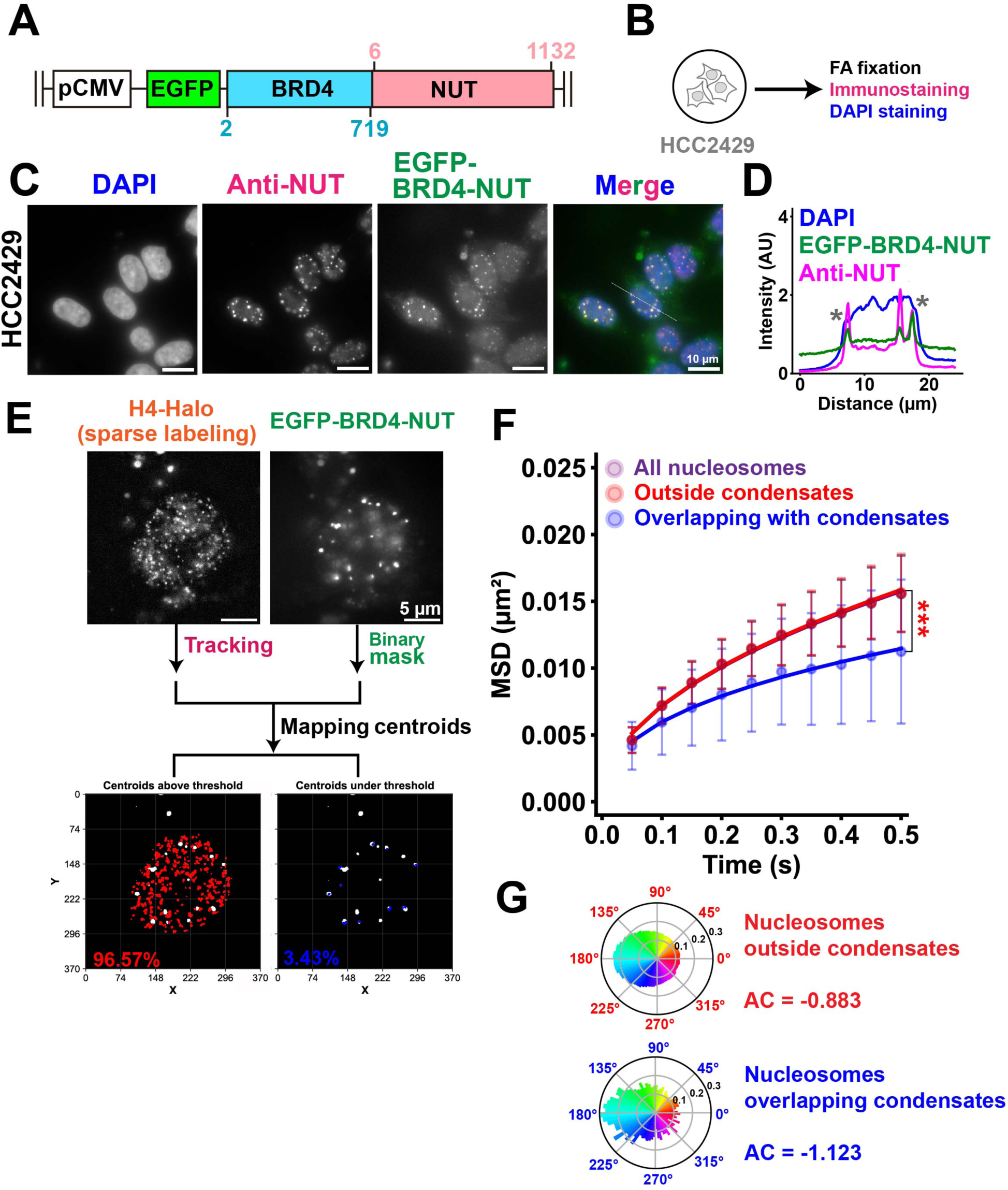
BRD4-NUT condensates constrain nucleosome motion in NC cells. **(A)** Schematic of the constitutive expression of EGFP-BRD4-NUT. **(B)** EGFP-BRD4-NUT condensates in HCC2429 cells do not require any induction or labeling. **(C)** Immunostaining against NUT (Anti-NUT, magenta) in HCC2429 cells with stable ectopic expression of EGFP-BRD4-NUT (green). Note that the weak foci observed in the nucleus are EGFP-BRD4-NUT condensates. Scale bars, 10 μm. Image intensities are optimized for visualization. **(D)** Line plot as derived from the thin white line crossing the nucleus in (C). Blue, DAPI (DNA); magenta, Anti-NUT; green, EGFP-BRD4-NUT. Note a significant overlap between EGFP-BRD4-NUT and Anti-NUT signal peaks. Asterisks (**\***) show the nuclear edge positions. **(E)** Simplified workflow in Fig. 4C optimized for HCC2429 cells. Top left: sparsely labeled H4-Halo image after background subtraction. Top right: EGFP-BRD4-NUT image in the same cell as top left. Bottom left: centroids of “outside” H4-Halo trajectories overlayed on the appropriate binary mask. % - their percentage from all trajectories. Bottom right: centroids of “overlapping” H4-Halo trajectories overlayed on the appropriate binary mask. % - their percentage from all trajectories. **(F)** MSD plots (± SD among cells) of “all” (purple), “outside” (red), and “overlapping” (blue) H4-Halo nucleosomes. For “all” and “outside”, *n* = 49 cells; for “overlapping”, *n* = 36 cells. ***, *p* = 5.1 × 10^−8^ for “overlapping” versus both “outside” and “all” by two-sided Kolmogorov-Smirnov test. **(G)** Measured angle distribution of “overlapping” and “outside” nucleosomes (5,543 angles for “overlapping”; 258,662 total number of angles for “outside”).

To test whether the BRD4-NUT condensates constrain euchromatic nucleosome motion in NC, we conducted euchromatin-specific single-nucleosome imaging in HCC2429 cells expressing H4-Halo (top left, Fig. 5E; right, Movie S5). As described for HeLa cells, we mapped the H4-Halo trajectory centroids onto binary masks created from the EGFP-BRD4-NUT condensate images (bottom, Fig. 5E). Again, nucleosome motions were constrained in the vicinity of the condensates (Fig. 5F, G). We conclude that the BRD4-NUT condensates locally constrain the movement of nucleosomes in NC, consistent with the result found in HeLa cells (Fig. 4F, G).

### BRD4-NUT crosslinks nucleosomes via its BRD4S region

We wondered how BRD4-NUT condensates locally constrain nucleosome movements. Because both chromatin binding and condensation of BRD4-NUT depend on the tandem bromodomains (BD1/2) of BRD4, which specifically recognize and bind to acetylated nucleosomes ^33,35,36^, we tested whether overexpression of BRD4 alone could affect nucleosome motion. Here, “BRD4” in BRD4-NUT refers to the short isoform of BRD4 (BRD4S), not the long isoform (BRD4L) ^36^. Unlike BRD4L (1,362 amino acids), BRD4S lacks the long intrinsically disordered C-terminal domain (CTD) and is only 722 amino acids in length ^53^. However, BRD4S retains both bromodomains (BD1/2) and is, in principle, capable of crosslinking and compacting acetylated nucleosomes ^35,54^.

We expressed inducible EGFP-BRD4S in HeLa cells already expressing H4-Halo (Fig. 6A). After a 24 hour induction, we observed that EGFP-BRD4S foci were smaller, more uniform in size, and much more abundant than EGFP-BRD4-NUT foci (lower, Fig. 6B, C). This is likely because BRD4S lacks NUT and cannot recruit p300.

**Fig. 6.**
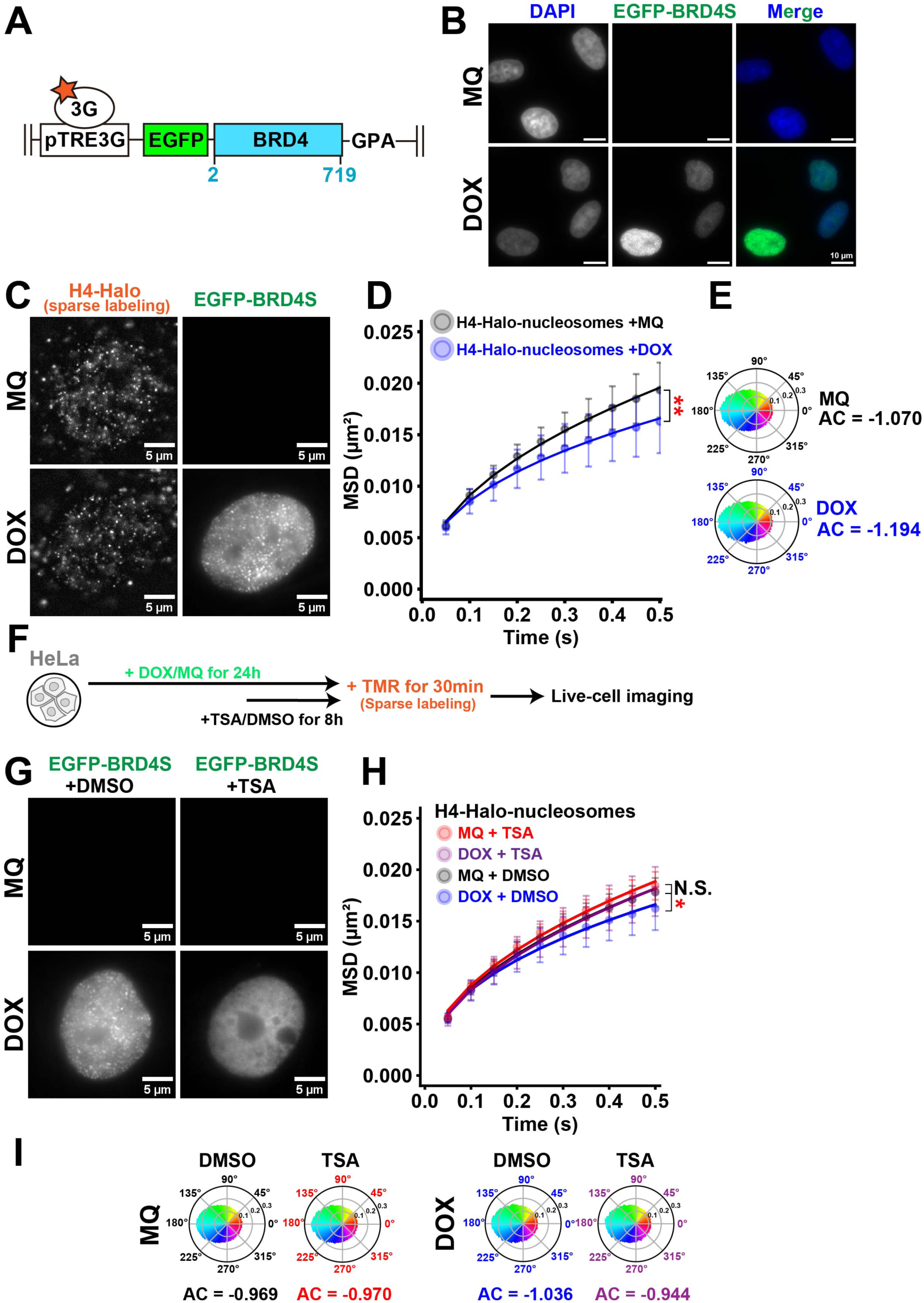
BRD4S forms condensates that also constrain nucleosome motion. **(A)** Schematic of the Tet-On 3G for doxycycline-inducible ectopic expression of EGFP-BRD4S. Note that BRD4S (722 aa) is almost identical to BRD4 (719 aa) in BRD4-NUT with an extra GPA tripeptide at the C-terminus. **(B)** HeLa cells were treated either with 0.1% Milli-Q (MQ) water or 1 µg/mL doxycycline (DOX) for 24 h before TMR labeling and FA fixation. Note that EGFP-BRD4S (BRD4S) foci are only seen in DOX treated and not in MQ treated cells. Scale bars, 10 μm. Image intensities are on the same scale. **(C)** Sparsely labeled H4-Halo image after background subtraction (left) and corresponding EGFP-BRD4S image (right) in MQ and DOX cells. Image intensities are optimized for visualization. **(D)** MSD plots (± SD among cells) of H4-Halo nucleosomes in MQ (black) and DOX (blue) cells. For each sample, *n* = 20 cells. ***, *p* = 4.0 × 10^−3^ for MQ versus DOX by two-sided Kolmogorov-Smirnov test. **(E)** Measured angle distribution of MQ and DOX nucleosomes (218,728 total number of angles for MQ; 165,652 angles for DOX). **(F)** Simplified protocol for 0.1% DMSO or 500 nM TSA treatment in the last 8 h during the 24-h incubation of cells in MQ or DOX. **(G)** Representative images of EGFP-BRD4S in cells treated with a combination of MQ/DOX and DMSO/TSA. Note that EGFP-BRD4S condensates observed as dots in DOX + DMSO are dispersed in DOX + TSA. Image intensities are on the same scale. **(H)** MSD plots (± SD among cells) of H4-Halo nucleosomes in MQ + DMSO (black, *n* = 21 cells), MQ + TSA (red, *n* = 20 cells), DOX + DMSO (blue, *n* = 19 cells), and DOX + TSA (purple, *n* = 23 cells) cells. *, *p* = 3.8 × 10^−2^ for MQ + DMSO versus DOX + DMSO; *p* = 1.9 × 10^−2^ for DOX + DMSO versus DOX + TSA; N.S. (not significant), *p* = 0.197 for MQ + DMSO versus DOX + TSA by two-sided Kolmogorov-Smirnov test. **(I)** Measured angle distribution of MQ + DMSO, MQ + TSA, DOX + DMSO, and DOX + TSA nucleosomes (236,522 total number of angles for MQ + DMSO; 271,029 angles for MQ + TSA; 134,973 angles for DOX + DMSO; 232,883 angles for DOX + TSA).

As it was too challenging to create binary masks of the very small and abundant EGFP-BRD4S foci, we analyzed the overall local motions of H4-Halo nucleosomes instead (Fig. 6D; right, Movie S6). Strikingly, euchromatic nucleosome motion was more constrained upon overexpression of EGFP-BRD4S (Fig. 6D, E), consistent with our findings from EGFP-BRD4-NUT (Figs. 4F-G and 5F-G). This result suggests that the ability of BRD4-NUT to crosslink acetylated nucleosomes likely originates from the tandem bromodomains of BRD4.

We then attempted to disperse the EGFP-BRD4S condensates with TSA (Fig. 6F), similar to what we did with EGFP-BRD4-NUT. Cells expressing EGFP-BRD4S were treated with TSA during the last 8 hours of the 24-hour doxycycline induction (Fig. 6F). The cells exhibited variable expression levels of EGFP-BRD4S. The condensates persisted in some cells with stronger expression but were dispersed in most cells, where the signals became evenly distributed throughout the nucleus (Fig. 6G). In cells with dispersed EGFP-BRD4S, euchromatic nucleosome motion returned to control levels (Fig. 6H, I; right, Movie S7). This demonstrates that BRD4S condensates functionally crosslink acetylated nucleosomes, revealing a possible mechanism by which the transcription machinery locally constrains nucleosome movements via BRD4-driven crosslinking during transcriptional activation (Fig. 7).

**Fig. 7.**
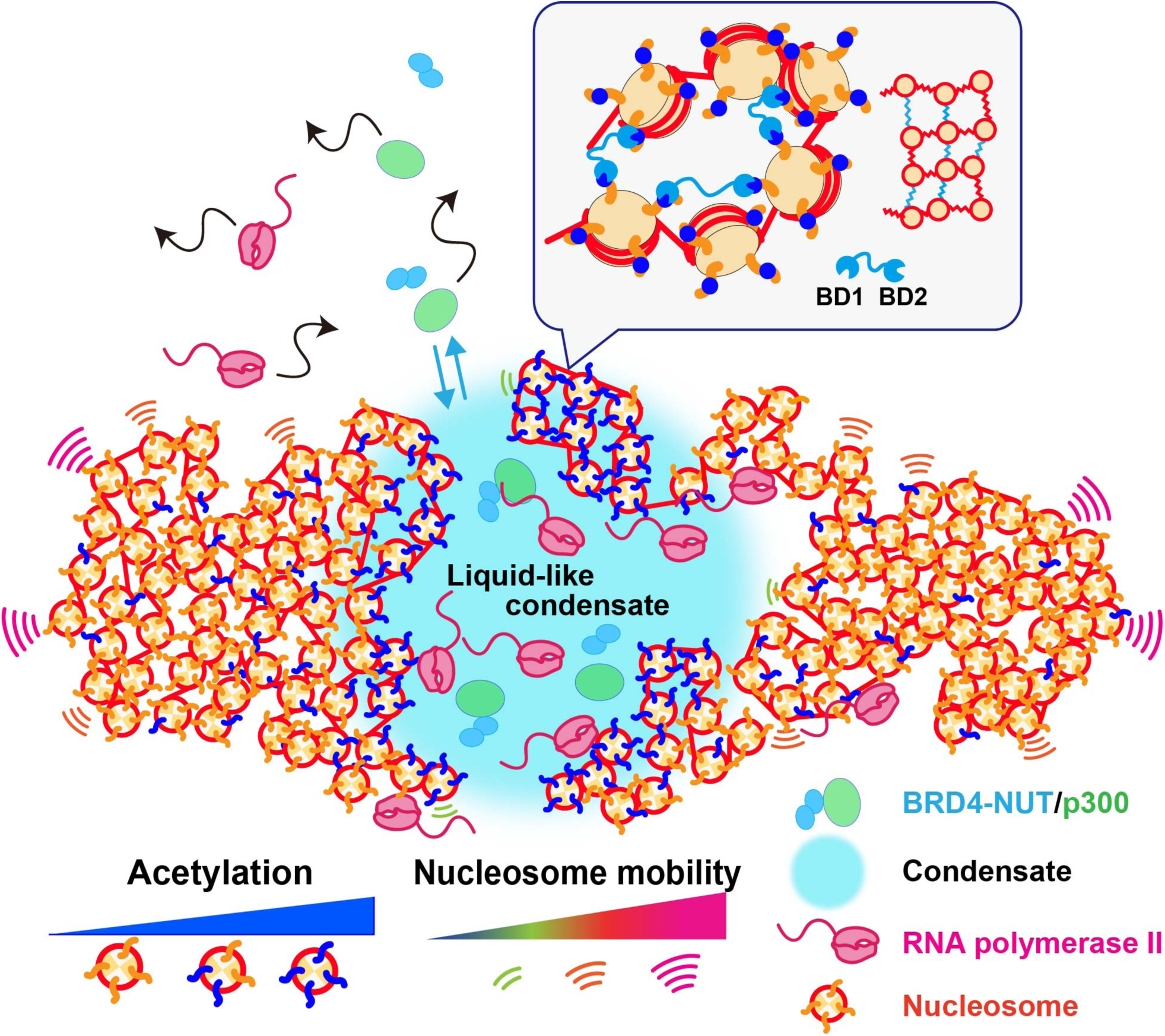
BRD4-NUT/p300 condensate locally constrains acetylated nucleosome motion. BRD4-NUT (blue ellipse) recruits and activates p300 (green ellipse) to form a complex. BRD4-NUT/p300 become much slower when trapped inside the viscous liquid-like condensate (large cyan circle). BRD4-NUT/p300 actively exchange between the freely diffusing and condensate populations (blue arrows). Active (phosphorylated) RNA Pol II complex (magenta cartoon) is also recruited to the condensate and drives aberrant gene transcription. The BRD4-NUT condensate locally constrains acetylated nucleosomes (nucleosomes with dark blue tails). In the inset, bromodomains BD1 and BD2 (found within BRD4) crosslink neighboring acetylated (dark blue spheres, left) nucleosomes to form a local gel-like structure that constrains chromatin (right).

We further confirmed that the activity of BRD4 bromodomains (BD1/2) is essential for nucleosome crosslinking using the BET bromodomain inhibitor JQ1 ^55^. JQ1 directly blocks BD1/2 at a 1:1 ratio (one JQ1 molecule per bromodomain) and prevents BRD4 from binding acetylated nucleosomes ^53^. When we treated cells with JQ1 during the final hour of the 24-hour doxycycline induction (Fig. S11A), we observed large spherical droplets of EGFP-BRD4S in the nucleus (bottom right, Fig. S11B; top right, Fig. S11G), which occasionally fused—consistent with their liquid-like physical properties ^53^. We inferred that JQ1 released all chromatin-bound EGFP-BRD4S condensates, allowing them to coalesce into large droplets. Interestingly, euchromatic nucleosomes became highly mobile (right, Movie S8)—even more so than in the control condition (Fig. S11C, D)—further supporting our conclusion that BD1/2 is involved in nucleosome crosslinking to constrain chromatin locally (Fig. 7).

Why did the mobility increase beyond control levels? Since we observed a marked reduction in the RPB1-S5P signal in these cells (Fig. S11E, F), we considered that transcription had already been inhibited by EGFP-BRD4S overexpression, but that the remaining condensates still restricted nucleosome motion. Nucleosomes became highly mobile only after being released from the condensates by JQ1. This interpretation was further supported by experiments using the CDK7 inhibitor THZ1 ^56^. THZ1 did not increase euchromatic nucleosome mobility any further in the background of EGFP-BRD4S overexpression and BET inhibition (Fig. S11H, I). Taken together, these findings strengthen our model that BRD4 condensates constrain local chromatin via BD1/2–acetylated nucleosome crosslinking (Fig. 7).

## Discussion

In this study, we employed single-molecule dual-color imaging to investigate the physical and functional properties of BRD4-NUT condensates in live human cells. Our results provide direct evidence that individual BRD4-NUT molecules diffuse within condensates like a viscous liquid. Notably, the MSD plots show that BRD4-NUT molecules in the condensates exhibit motion similar to that of molecules in the nucleoli ^57^, which are highly crowded and known to be viscous liquid droplets ^21,58^. Strikingly, these viscous liquid-like condensates locally constrain nucleosome motion in euchromatic regions (Fig. 7), which correspond to Hi-C A-compartments enriched in active histone marks (Fig. S7) ^38^. These findings fill a critical gap in our understanding of how transcriptional condensates formed via LLPS can modulate chromatin dynamics, particularly in the context of active gene regulation. Our work is also compatible with the classical transcription factory model, where stable clusters of RNA Pol II work as transcription factories and immobilize chromatin to be transcribed ^17,59–61^, and provides a molecular basis to the model.

For this study, we developed three imaging approaches to investigate the molecular assembly of condensates and their interplay with chromatin dynamics. First, by labeling Halo-BRD4-NUT sparsely with JF646 and densely with TMR, we simultaneously tracked single molecules inside the condensates and the movement of entire condensate foci (Fig. 2). This allowed us to observe the slow, but dynamic behavior of Halo-BRD4-NUT within these condensates. Second, we performed dual-color labeling and simultaneous imaging of Halo-JF646-BRD4-NUT and Halo-TMR-BRD4-NUT (Fig. 3). The restricted localization of BRD4-NUT molecules to discrete nuclear condensates provided a significant advantage in identifying closely positioned molecule pairs. We analyzed the two-point MSD between these pairs, as this metric is less affected by translational or rotational movements of the condensates. The resulting linear two-point MSD plots provided direct evidence that BRD4-NUT condensates in cells exhibit viscous liquid-like behavior, because linear MSD is a characteristic of normal (or free) diffusion. Finally, using euchromatin-specific single-nucleosome imaging and tracking, we developed a method to analyze nucleosome motion around the condensates by mapping nucleosome trajectories onto binary masks generated from condensate images (Fig. 4). Ectopically expressed H4-Halo is preferentially incorporated into euchromatin-associated nucleosomes with active chromatin marks (Fig. S7) ^38^, enabling us to focus on dynamic processes primarily occurring in transcriptionally active euchromatin. Our findings reveal that nucleosome motion is indeed locally constrained around BRD4-NUT condensates (Fig. 7).

These single-molecule-based methods advance our understanding of the molecular assembly of biomolecular condensates, offering an alternative to FRAP, which does not directly determine whether these structures exhibit liquid-like properties ^21,24^. Combining these approaches with further perturbations—such as rapid protein depletion ^62^, optogenetic control ^63,64^, or other forms of chromatin manipulation—may further elucidate how phase-separated domains interface with the physical properties of the genome.

Finally, one of the most striking findings of our study is that liquid-like BRD4-NUT condensates physically restrict the motion of nucleosomes located in their vicinity. This effect was observed both in engineered HeLa cells and in NC-derived HCC2429 cells, supporting its biological relevance. While synthetic condensates have been shown to interact with chromatin and probe its physical properties ^64^, to our knowledge, this is the first demonstration that native condensates can actively modulate the physical state of chromatin in living cells. Furthermore, we found that overexpression of BRD4S alone—lacking the NUT region—was sufficient to reduce nucleosome mobility, implying that the NUT region does not play a major role in constraining nucleosomes. But, as it binds to and activates p300 ^35^, the NUT region serves a regulatory function by modulating condensate size in NC-derived cells. The activity of BRD4S was abolished by the BET inhibitor JQ1 or by hyperacetylation via TSA treatment, demonstrating that the interaction between the bromodomains (BD1/2) of BRD4S and acetylated nucleosomes is essential for neighboring nucleosome crosslinking and local constraint (Fig. 7). Thus, BRD4-driven condensates are not passive assemblies but actively modulate chromatin mechanics through BD1/2-mediated binding. Our results suggest a general mechanism by which LLPS-based condensates can modulate chromatin dynamics via physical interactions with nucleosomes. This adds a new dimension to the functional repertoire of transcriptional condensates, which have largely been considered as compartments that concentrate some components while excluding others to promote efficient biochemical reactions.

## Materials and methods

### Construction of expression plasmids

The plasmid for the doxycycline-inducible expression of EGFP-BRD4-NUT to be integrated into the AAVS1 safe harbor locus on chromosome 19 via CRISPR/Cas9 genome editing was made first. The pCMV-EGFP-BRD4-NUT-C1 expression plasmid was described in ^33,35^. The EGFP-BRD4-NUT coding sequence was amplified to replace the KRAB-dCas9 (CRISPRi) region from pAAVS1-NDi-CRISPRi (Gen 2, Addgene #73498). The HaloTag version of the plasmid was constructed similarly by combining the identical vector, BRD4-NUT, and HaloTag sequences. The analogous plasmid for EGFP-BRD4S was made by amplifying the construct excluding the NUT sequence with added 5’-GGACCAGCA-3’ nucleotides (corresponding to Gly-Pro-Ala amino acids at the C-terminus of the BRD4S sequence). To make the plasmid for constitutive expression of EGFP-BRD4-NUT, the EGFP-BRD4-NUT sequence was amplified and then inserted into the multiple cloning site (MCS) on the PB-CMV-MCS-EF1-Puro plasmid (PB510B-1, System Biosciences).

### Establishment of stable cell lines

HeLa cells containing the flippase recognition target (FRT) site in their genome, HeLa FRT-bla ^65^, and HeLa cells constitutively expressing histone H4-HaloTag under the EF1 promoter, HeLa H4-Halo, were used. HeLa cells were cultured in Dulbecco’s Modified Eagle medium (DMEM, D5796-500ML; Sigma-Aldrich) supplemented with 10% fetal bovine serum (FBS, ABB213138; HyClone) at 37°C in 5% CO_2_. HCC2429 cells ^33^were cultured in RPMI1640 medium (189-02025, Fujifilm) supplemented with 10% FBS at 37°C in 5% CO_2_.

To establish HeLa cells with doxycycline-inducible ectopic expression of EGFP-BRD4-NUT, Halo-BRD4-NUT, or EGFP-BRD4S, the corresponding plasmids were co-transfected with pX330-AAVS1 (Addgene #227272) using the Effectene Transfection Reagent kit (301425; QIAGEN) in a 35-mm dish. Two days following the transfection, the cell populations were expanded into 100-mm dishes with medium containing 500 μg/mL G418 (ALX-380-013; Enzo) for positive selection. The medium was refreshed every 3-4 days. Clones were isolated after ∼2 weeks of growth. The PiggyBac system was used for the stable ectopic expression of EGFP-BRD4-NUT or H4-HaloTag in HCC2429 cells as follows. PB-CMV-EGFP-BRD4-NUT-EF1-Puro was co-transfected with the PiggyBac transposase expression plasmid, pCMV-hyPBase ^66^, using the Effectene Transfection Reagent kit in a 12-well plate and selected for with 1 μg/mL puromycin dihydrochloride (P8833; Sigma Aldrich). PB-EF1-H4-Halo-IRES-Neo ^38^ was co-transfected with pCMV-hyPBase using the Effectene Transfection Reagent kit in a 12-well plate and selected for with 800 μg/mL G418.

### Expression and localization of H4-HaloTag, BRD4-NUT, and RFP-p300 in cells

To assess the localization of H4-HaloTag in HeLa and HCC2429 cells, cells were seeded onto 1 mg/mL poly-L-lysine-coated (P1524-500MG; Sigma-Aldrich) coverslips (C018001; Matsunami). The HaloTag was labeled with an excess amount of ligand, 5 nM tetramethylrhodamine (TMR, 8251; Promega), for 30 min at 37°C in 5% CO_2_. The rest of the procedures were conducted at room temperature.

The localization of Halo-BRD4-NUT in HeLa cells was assessed following a 24-h induction with culture medium (DMEM supplemented with 10% FBS) containing 1 μg/mL doxycycline (or 0.1% Milli-Q water for control). EGFP-BRD4-NUT expression was similarly induced and imaged in HeLa cells as described below.

For the transient expression of both RFP-p300 and EGFP-BRD4-NUT in HeLa cells, pCMV-TagRFP-p300-C1 ^35^ and pAAVS1-NDi-EGFP-BRD4-NUT were co-transfected using the Effectene Transfection Reagent kit in a 35-mm dish. The following day, the culture medium was exchanged for DMEM with 10% FBS containing 1 µg/mL doxycycline. The localizations of RFP-p300 and EGFP-BRD4-NUT were assessed 24 h after doxycycline addition.

Cells grown on coverslips were briefly rinsed with 1×PBS and then fixed with 1.85% formaldehyde (064-00406; Wako) in 1×PBS for 15 min. Excess molecular crosslinking and fixation were quenched with 50 mM glycine (077-00735; Wako) in 1×HMK (20 mM HEPES (pH 7.5) with 1 mM MgCl_2_ and 100 mM KCl) for 5 min. The cells were then permeabilized with 0.5% Triton X-100 (T-9284; Sigma-Aldrich) in 1×HMK for 5 min. Wherever applicable, indirect immunocytochemistry (see section below) was conducted prior to DNA staining. After briefly rinsing with 1×HMK, DNA was stained with 0.5 µg/mL 4’,6-diamidino-2-phenylindole (DAPI, 10236276001; Roche) in 1×HMK for 5 min. Finally, after rinsing with 1×HMK twice, the coverslips were mounted onto micro slide glass slides (S011120; Matsunami) in *p*-phenylenediamine (PPDI; 20 mM Hepes (pH 7.4), 1 mM MgCl_2_, 100 mM KCl, 78% glycerol, and 1 mg/mL paraphenylene diamine (695106-1G; Sigma-Aldrich)) and sealed with nail polish (Kate Top Coat N 01, Kanebo).

Z-stack images (every 0.4 µm along the Z axis, 20-25 sections in total) of the cells were taken using a DeltaVision Ultra (GE Healthcare) epifluorescence microscope equipped with an Olympus PlanApoN 60× NA 1.42 objective, sCMOS camera, InsightSSI light (∼50 mW), and the four-color standard filter set. DeltaVision image acquisition software, SoftWorx 7.X, was used to acquire images in all cases. Image fusion, projection through the entire nucleus and quantification from the projections were done using Fiji (ImageJ).

To quantify the intranuclear fluorescence signal intensity, DAPI-stained nuclear regions were segmented with Fiji based on the Otsu thresholding method. The mean nuclear pixel intensities (16-bit arbitrary units, A.U.) were measured inside the segmented regions. Line plots were made by measuring the signal intensities along ∼24 µm lines crossing the nuclei or ∼1.5 µm lines crossing the fluorescent dots, and the values were normalized and adapted for visualization.

### Indirect immunocytochemistry

Upon permeabilization with 0.5% TritonX-100 in 1×HMK (see previous section), the cells were incubated in 10% normal goat serum (NGS; 143-06561, Wako) in 1×HMK for 30 min. The cells were then incubated with one of the following primary antibodies: 1) rabbit anti-RPB1 S5P (1:1000; ab5131, Abcam), 2) rabbit anti-acetylated histone H3 (1:1000; 06-599, Millipore), 3) rabbit anti-NUT (1:1000; 3625S, Cell Signaling) diluted in 1% NGS in HMK for 1 h. After being washed with HMK four times, the cells were incubated in the secondary antibody, goat anti-rabbit IgG Alexa Fluor 647 (1:1000; A212445, Abcam), in 1% NGS in HMK for 1 h followed by a wash with HMK four times. Then, DNA staining was performed as previously described.

### Pharmacological treatments

- 2% FA: For single-molecule imaging in FA-fixed cells, the cells were incubated in 2% FA in 1×PBS at RT for 30 min, washed with 1×PBS two times, and kept in 1×PBS at 4°C for >4 h. The cells were returned to the culture medium at 37°C in 5% CO_2_ before imaging.
- 500 nM TSA: The cells were incubated in the culture medium containing 1 μg/mL doxycycline (631311, BD) or 0.1% Milli-Q water for 24 h. In the last 8 h of this treatment, the cells were incubated in 500 nM TSA (203-17561, Wako) or 0.1% DMSO (D2650, Sigma Aldrich).
- 1 μM JQ1: The cells were incubated in the culture medium containing 1 μg/mL doxycycline or 0.1% Milli-Q water for 24 h. In the last 1 h of this treatment, the cells were incubated in 1 μM (+)-JQ1 (SML1524, Sigma Aldrich) or 0.1% Milli-Q water.
- 1 μM THZ1: The cells were incubated in the culture medium containing 1 μg/mL doxycycline or 0.1% Milli-Q water for 24 h. In the last 2 h of this treatment, the cells were incubated in 1 μM (+)-JQ1 or 0.1% Milli-Q water and 1 μM THZ1 hydrochloride (CS3168, Funakoshi) or 0.1% DMSO.

### Single-molecule imaging microscopy

To assess the local nucleosome behavior of H4-Halo in living HeLa and HCC2429 cells, the cells were seeded onto 1 mg/mL poly-L-lysine-coated glass-base dishes (3970-035; Iwaki). The HaloTag was sparsely labeled with 50 pM TMR in phenol red-free DMEM (HeLa; 21063-029, Thermo Fisher Scientific) or RPMI1640 (11835030, Thermo Fischer Scientific) with 10% FBS for 20 min at 37°C in 5% CO_2_. The cells were then washed with Hank’s balanced salt solution (1×HBSS, H1387; Sigma Aldrich) three times and kept in phenol red-free DMEM or RPMI1640 with 10% FBS at 37°C in 5% CO_2_ using an INU-TIZ-F1 (Tokai Hit) live cell chamber and GM-8000 digital gas mixer (Tokai Hit) during imaging. Single H4-HaloTag-TMR nucleosomes were visualized using a Nikon Eclipse Ti inverted microscope with a 100-mW Sapphire 561-nm laser (Coherent) and sCMOS ORCA-BT camera (Hamamatsu Photonics). TMR was excited by the 561-nm laser through the Nikon 100× PlanApo TIRF NA 1.49 objective and detected at 575–710 nm. Oblique illumination with a thin optical sheet achieved high signal-to-noise ratio for single-nucleosome imaging ^44–46^. Sequential image frames (*n* = 300) were captured using NIS elements software (AR v5.30.03 64bit, Nikon) at 50 ms of continuous laser exposure.

To assess the single-molecule behavior of Halo-BRD4-NUT in living HeLa cells, the analogous procedure was followed with several key differences. HaloTag was sparsely labeled with 50 pM JF646 (GA1120, Promega) in phenol red-free DMEM with 10% FBS for 40 min at 37°C in 5% CO_2_. For dense HaloTag labeling, the cells were incubated in 5 nM TMR in the last 20 min of JF646 treatment. Single Halo-JF646-BRD4-NUT and Halo-TMR-BRD4-NUT condensates were visualized with 40-mW 640-nm and 100-mW 561-nm lasers, respectively (Coherent). JF646 was excited by the 640-nm laser and detected at 676-786 nm, whereas TMR was excited by the 561-nm laser and detected at 582–626 nm. Sequential image frames, 500 (or 300) for each channel were captured using NIS elements software at a frame rate of 10 ms (or 50 ms).

To achieve dual-color single-molecule imaging, the TMR concentration was reduced to 10 pM. Halo-JF646-BRD4-NUT and Halo-TMR-BRD4-NUT single dots were simultaneously visualized using a beam splitter system, W-VIEW GEMINI (Hamamatsu Photonics).

### Single-molecule and condensate tracking and analysis

Image processing and preparation for single-molecule tracking were performed using Fiji. The background signal was subtracted using the rolling ball method (radius = 50 pixels). The region corresponding to the cell nucleus was manually extracted using the “Polygon selections” method. The centroids of detected fluorescent dots in each image were determined, and their trajectories were calculated using u-track (H4-Halo and Halo-JF646-BRD4-NUT at 50 ms; MATLAB package ^67^) or TrackMate (Halo-JF646-BRD4-NUT at 10 ms; Fiji package ^47^). For TrackMate, the LoG detector was used for detection of single molecule dots, and the linking distance was set to 2 pixels. The u-track and TrackMate trajectories were then used to calculate the displacement distributions and the MSDs of fluorescent dots in Python. The originally calculated 2D MSD values were multiplied by 1.5 (4 to 6 *D*t) to obtain 3D MSD values. MSD plots with appropriate statistical analyses between various phenotypes were made using Python.

To categorize whether a given H4-Halo or Halo-JF646-BRD4-NUT trajectory belongs to a condensate of EGFP/Halo-BRD4-NUT or not, the EGFP/Halo-BRD4-NUT condensate image was passed through the “Bandpass filter” with the minimum of 3 pixels and the maximum of 20 pixels. Then, the obtained image was automatically binarized using the Otsu or MaxEntropy method. If the centroid of the H4-Halo trajectory fell in the white region determined by the binary mask, the entire trajectory was categorized as “overlapping with the condensate”. Otherwise, it was categorized as “outside the condensate”. MSDs were separately calculated for each category of H4-Halo and Halo-JF646-BRD4-NUT trajectories.

Image processing and preparation for Halo-TMR-BRD4-NUT condensate tracking were performed using Fiji. The first 100 frames of condensate images were passed through the “Bandpass filter” with the minimum of 3 pixels and the maximum of 20 pixels. The obtained movies were automatically binarized using the Otsu method at each frame. The irregularly shaped white regions in each image were detected using the Thresholding detector of TrackMate, and their centroids were tracked similarly to single molecules (linking distance = 2 pixels). To estimate the condensate dimensions for two-point MSD analysis, the “Bandpass filter” for binarization was set with the minimum of 3 pixels and the maximum of 6 pixels to obtain crisper condensate outlines.

For angle distribution analysis ^48,52^, the consecutive tracked positions {(*x*_0_, *y*_0_), (*x*_1_, *y*_1_), ⋯, (*x_n_*, *y_n_*), ⋯} of single molecules and condensates on the (*x*, *y*) plane were first converted into a set of displacement vectors Δ***r***_n_ = (*x*_n + 1_ − *x*_n_, *y*_n + 1_ − *y*_n_)*^t^*. Then, the angle between two vectors Δ***r***_n_ and Δ***r***_n + 1_ was calculated for every trajectory in our experiments, and the normalized polar histogram was plotted in Python. The angle distribution was normalized by 2π, and the magnitudes corresponded to the probability density. The color spectrum indicates each angle in a circular range from 0° to 360°.

### Two-point MSD analysis

Image processing, dot detection, and tracking of the dual-color single-molecule tracking images were carried out using TrackMate (LoG detector, linking distance = 2 pixels) as described above. The calculated position determination accuracy through the beam splitter was 16.3 nm (TMR particles) and 14.3 nm (JF646 particles) (Fig. S6C and E). Positions of spots labeled by TMR and JF646 colors were corrected by second-order polynomial transformation. Parameters of the transformation were estimated based on the 0.1-μm TetraSpeck bead (T7279, Thermo Fisher Scientific) imaging and calculations were performed using the Python library Scikit image. To analyze the movements of two molecules labeled with two colors (TMR and JF646), we picked closely located (<300 nm) pairs of TMR and JF646 trajectories from each image movie. Around 3,500 pairs of the dots were obtained from a single experiment. The two-point MSD of the fluorescent dots was calculated as described in ^7,50^. In our single-molecule tracking, trajectories of TMR- and JF646-dots on the (*x*, *y)* plane, 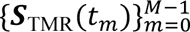 and 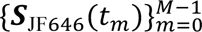, were simultaneously acquired, where the time interval was Δ*t* = 0.01 s and *t_m_* = *m*Δ*t* (*m* = 0,1,2, ⋯, *M* − 1). To evaluate dynamic fluctuations between two points, we dealt with the relative vector between two trajectories, ***Q***(*t_m_*) = ***S***_TMR_(*t_m_*) − ***S***_JF646_(*t_m_*). Then, we calculated the two-point MSD for the lag time *t*_*_ = *n*Δ*t* by

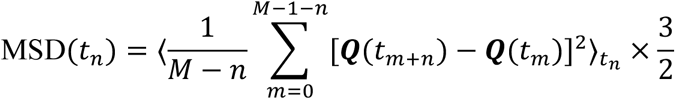

where 〈·〉*t*_n_ represents the ensemble average for trajectories at the lag time *t*_n_ and the coefficient ^3^/_2_ is a correction factor for conversion from 2D to 3D values.

### Cellular protein preparation and immunoblotting

HeLa cells were lysed in hot Laemmli buffer ^68^ containing 0.1 M dithiothreitol (DTT; 040-29223, Wako). The resulting lysate was incubated at 95°C for 10 min to denature proteins and then sonicated. The proteins from the lysate were resolved on an 8% polyacrylamide SDS-gel at 150 V for 2 h (Fig. S3C, D and Fig. S11E, F) or 10% gel at 150 V for 1 h (Fig. S3A, B). Gels were transferred to a 0.45-μm polyvinylidene difluoride (PVDF) Immobilon-P membrane (IPVH00010, Merck) at a constant current of 110 mA for 1 h using the semi-dry method (ATTO; 25 mM tris, 5% methanol). The membranes were blocked in 5% bovine serum albumin (Figs. S3A, B and S11E, F; BAC65-1150, Equitech-Bio) or 5% skim milk (Fig. S3C, D case, 190-12865, Wako) in phosphate-buffered saline (PBS) or Tris-buffered saline (TBS) with 0.1% Tween 20 for 1 h at room temperature. The membrane-bound proteins were probed with anti-H4 (1:4000 dilution; ab10158, Abcam), anti-HaloTag (1:1000 dilution; G9211, Promega), anti-lamin A/C (1:1000 dilution; sc-7292, Santa Cruz), anti-GFP (1:5000; 598, MBL), anti-BRD4 (1:1000 dilution; ab128874, Abcam), anti-NUT (1:1000 dilution; 3625S, Cell Signaling), anti-RPB1 S5P (1:1000 dilution; ab5131, Abcam), or anti-RPB2 (1:1000 dilution; sc-166803, Santa Cruz) antibodies, followed by the appropriate secondary antibody: anti-rabbit (1:5000 dilution; 170-6515, Bio-Rad) or anti-mouse (1:5000 dilution; 170-6516, Bio-Rad) horseradish peroxidase (HRP)–conjugated goat antibody. Chemiluminescence reactions were used (WBKLS0100, Millipore) and detected by EZ-Capture MG (AE-9300H-CSP, ATTO).

## Acknowledgments

We are grateful to Dr. K.M. Marshall, Dr. S. Shinkai, and Mr. Y. Nagata for critical reading of this manuscript and Ms. S. Tamura for figure illustration. We thank Dr. H. Kimura for providing his antibodies, Dr. S. Iida, Dr. K. Hibino, and Maeshima laboratory members for their helpful discussions and support. This work was supported by JSPS grants JP24H0061 (K.Maeshima), JP22H05606 (S.I.), JP23K17398 (K.Maeshima, S.I.), 22H04925 (PAGS)(K.Maeshima), Takeda Science Foundation (K.Maeshima). A.S. is a MEXT Scholar. K.Minami was a SOKENDAI Special Researcher (JST SPRING JPMJSP2104) and a JSPS Fellow (JP23KJ0998). M.A.S. is a JSPS Fellow (24KJ1161).

## Author contributions

A.S. and K.Maeshima designed the project. A.S. generated most of the cells, partly with the help of S.I. A.S. performed single-nucleosome imaging, tracking, and analysis. A.S. performed the cell biology experiments. K.Minami contributed to the analysis of single BRD4-NUT and its condensates. M.A.S. contributed to two-point MSD. A.S. and K.Maeshima wrote the manuscript with input from all the authors.

## Data and materials availability

The sequencing data have been deposited with links to BioProject accession numbers PRJDB17378 (H2B-HaloTag pull-down) and PRJDB17379 (H4-HaloTag pull-down) in the DDBJ BioProject database ^38^. The scripts for track sorting and angle distribution analysis are available at https://doi.org/10.5281/zenodo.15812848. Other data are available in the main text or the supplementary materials.

---

**Fig. S1.**
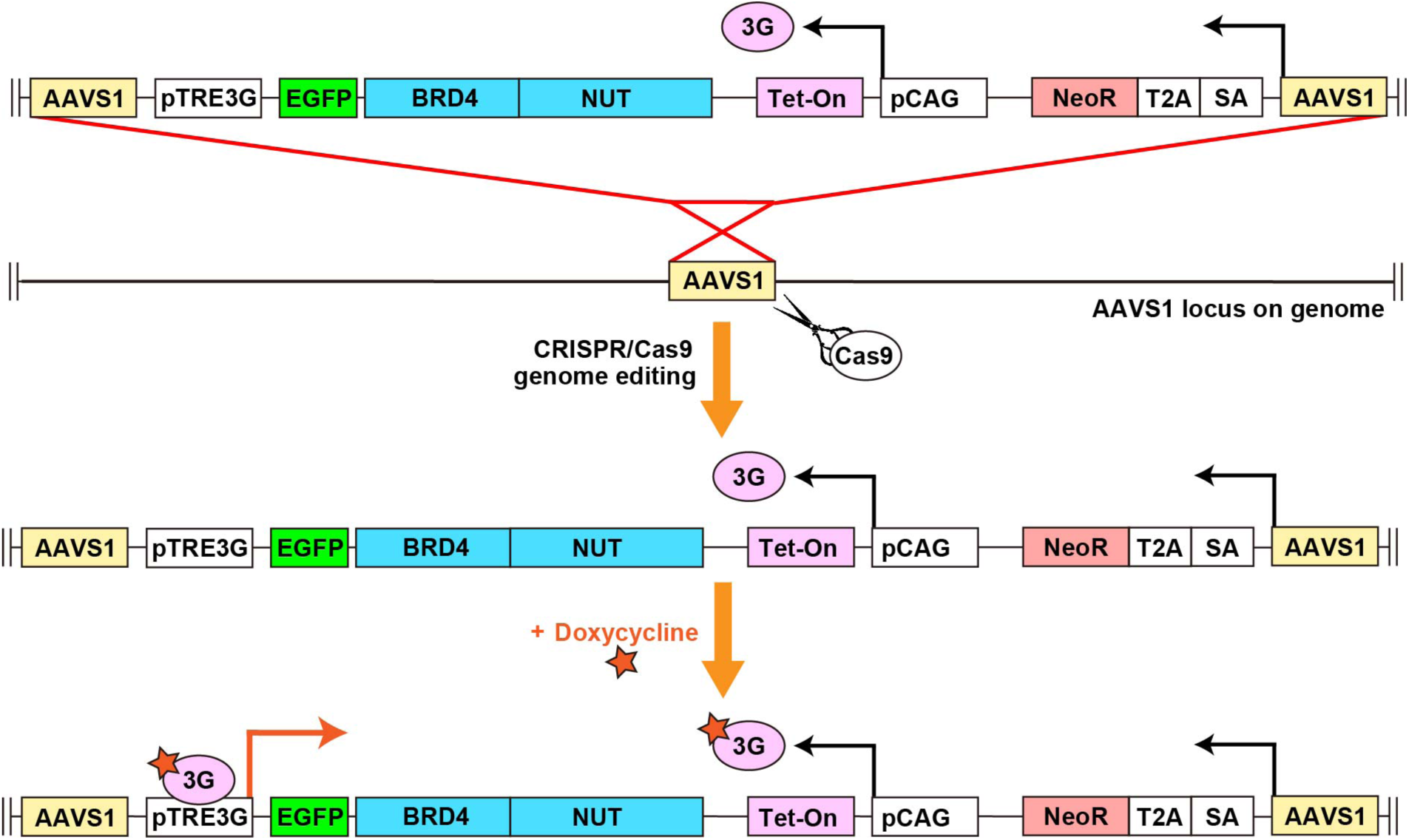
Schematic of the Tet-On 3G for doxycycline-inducible ectopic expression of EGFP-BRD4-NUT. Note that the construct is inserted into the AAVS1 safe harbor locus via CRISPR/Cas9 genome editing, and the cells are positively selected with G418 antibiotic. An analogous strategy was used for the inducible expression of Halo-BRD4-NUT and EGFP-BRD4S.

**Fig. S2.**
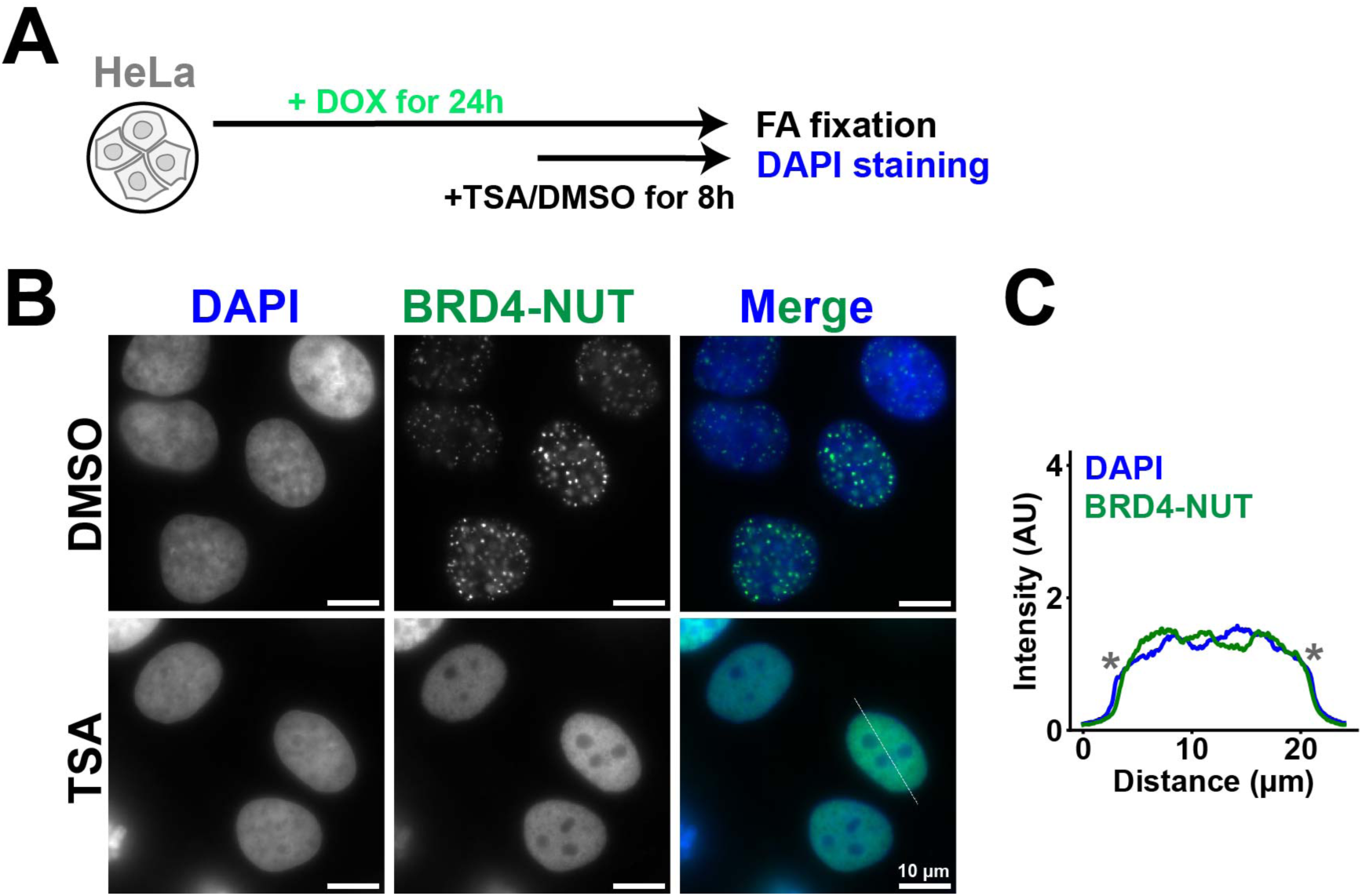
BRD4-NUT condensates are dispersed with TSA treatment. **(A)** Simplified protocol for DMSO or TSA treatment in the last 8 h during the 24-h incubation of cells in 1 µg/mL doxycycline (DOX). **(B)** EGFP-BRD4-NUT images in HeLa cells treated either with DMSO (0.1%) or TSA (500 nM) in the last 8 h of the 24-h induction with 1 μg/mL doxycycline before FA fixation. Scale bars, 10 μm. Image intensities are not on the same scale, but were optimized for visualization. **(C)** Line plot as derived from the thin white line crossing the nucleus from (B). Blue, DAPI (DNA); green, EGFP-BRD4-NUT. Note a weak anticorrelation between EGFP-BRD4-NUT and DAPI signal peaks. Asterisks (**\***) show the nuclear edge positions.

**Fig. S3.**
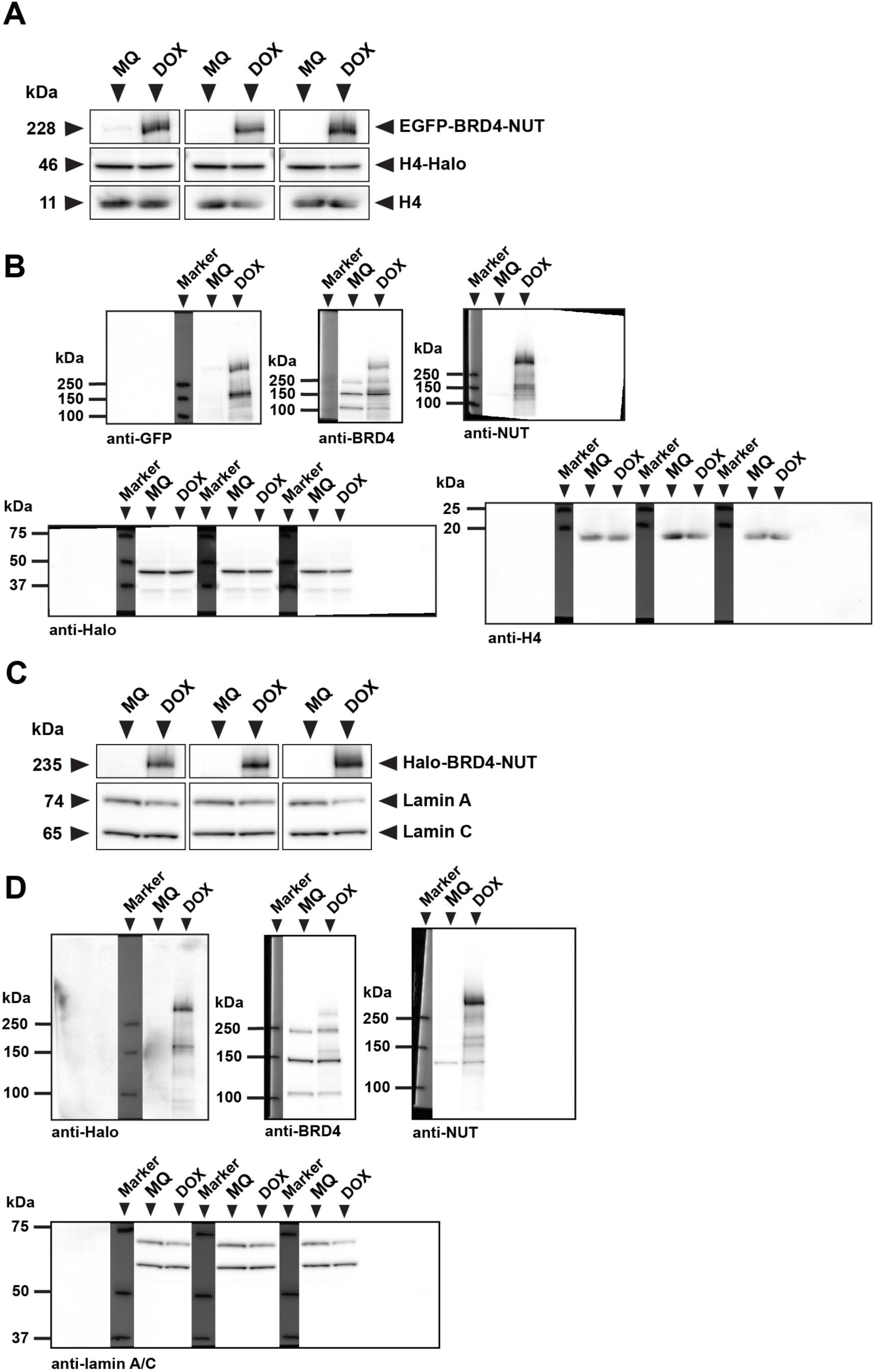
Immunoblots of EGFP- and Halo-BRD4-NUT from HeLa cell lysates. **(A)** Immunoblotting against EGFP-BRD4-NUT, histone H4, and H4-Halo in MQ or DOX treated HeLa cells. EGFP-BRD4-NUT was detected with: anti-GFP, anti-BRD4, and anti-NUT antibodies. Note that EGFP-BRD4-NUT band appears only in lysates from DOX treated cells. **(B)** Uncropped images of the PVDF membranes used for immunoblotting in (A). Membranes were incubated with anti-GFP, anti-BRD4, anti-NUT, anti-Halo, and anti-H4 antibodies. **(C)** Immunoblotting against Halo-BRD4-NUT and lamin A/C in lysates from MQ and DOX treated HeLa cells. Halo-BRD4-NUT was detected with: anti-Halo, anti-BRD4, and anti-NUT antibodies. Note that the Halo-BRD4-NUT band appears only in lysates from DOX treated cells. **(D)** Uncropped images of the PVDF membranes used for immunoblotting in (C). Membranes were incubated with anti-Halo, anti-BRD4, anti-NUT, and anti-lamin A/C antibodies.

**Fig. S4.**
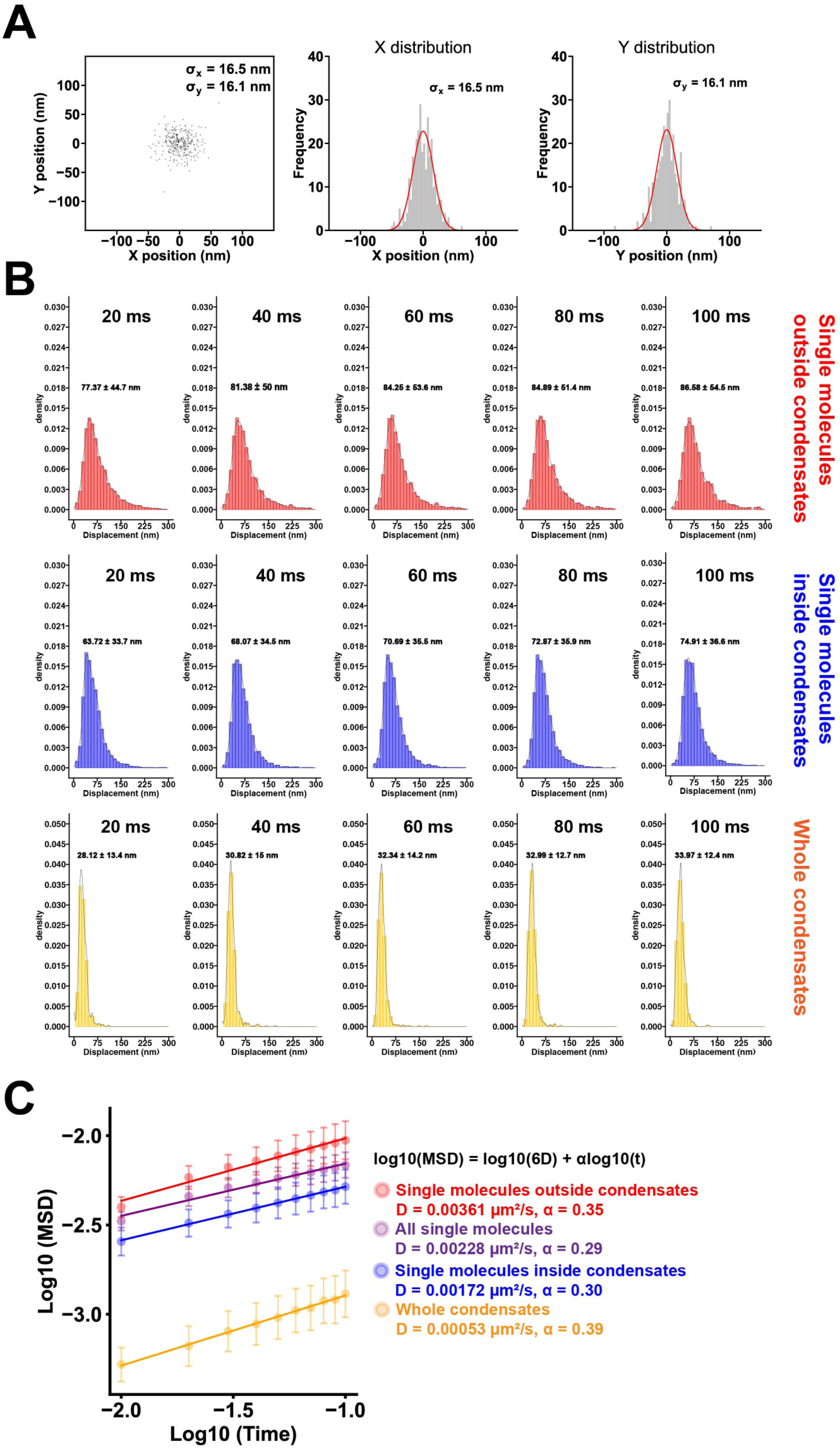
Position determination accuracy and displacement histograms of Halo-BRD4-NUT dots. **(A)** Position determination accuracy of single Halo-BRD4-NUT molecules. Distribution of single-molecule displacements from the centroids in the (*x, y*)-plane over the 10-ms interval, *n* = 10 molecules. SD*_x_* and SD*_y_* of fitted Gaussian functions were 16.5 nm and 16.1 nm, respectively. **(B)** Displacement distributions of Halo-BRD4-NUT (*n* = 20 cells) for 20, 40, 60, 80, and 100 ms. Means ± SD of displacement are indicated in the center. **(C)** Log-log plot of the MSD data in Fig. 2E. The indicated lines were fitted using the data from 0.01 to 0.1 s. The formula shown in black was used to calculate *D* and α values for each plot.

**Fig. S5.**
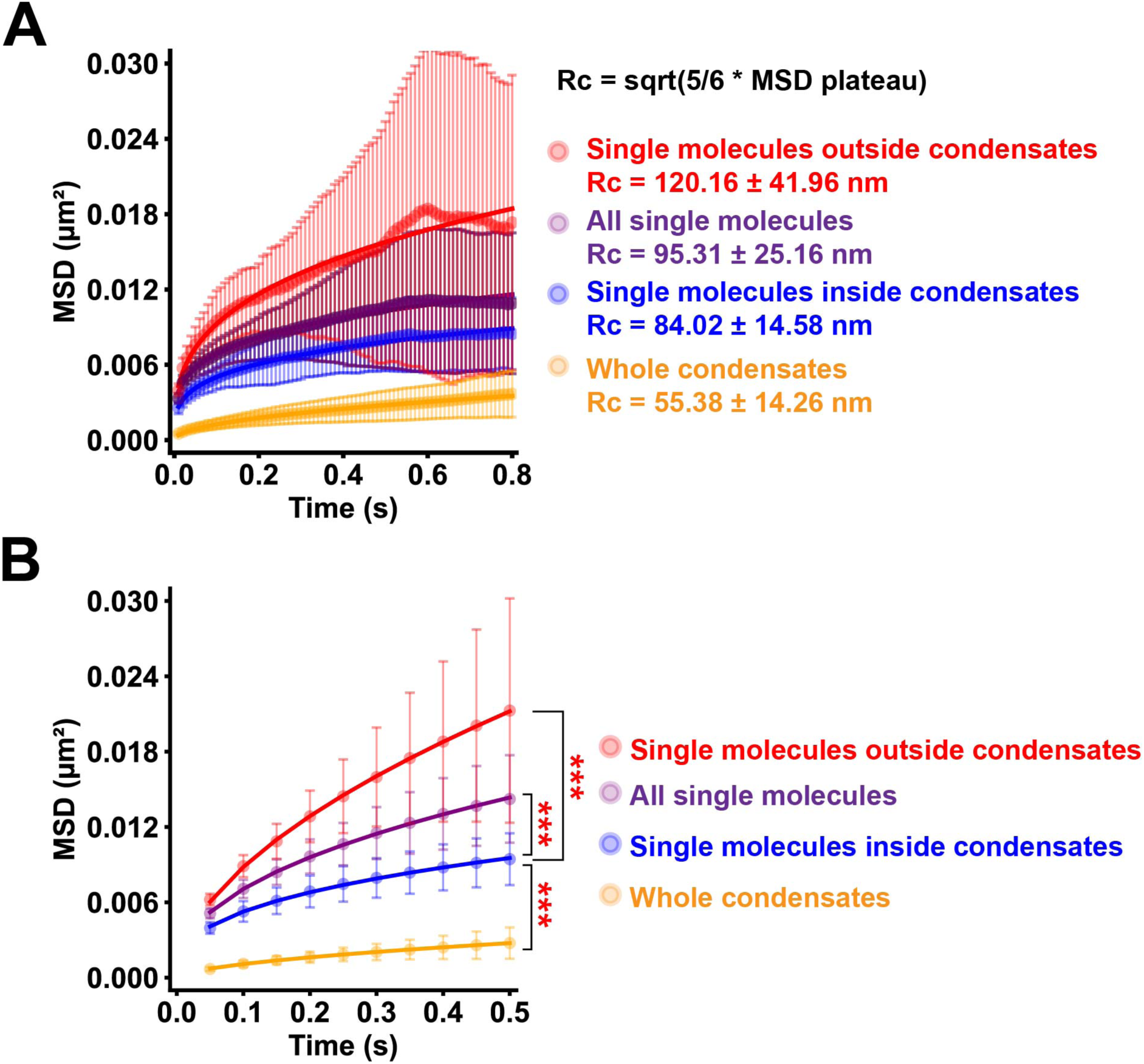
Single- and dual-color single-molecule imaging of BRD4-NUT at 50 ms/frame. **(A)** MSD plots (± SD among cells) of Halo-BRD4-NUT “all” (purple), “inside” (blue), and “outside” (red) JF646 labeled single molecules, and TMR labeled whole condensates (orange) over a longer time range (0.01 to 0.8 s). MSD plateau was achieved at 0.8 s. The formula shown in black was used to calculate *R*_c_ values (mean ± SD) for each plot. **(B)** MSD plots (± SD among cells) of Halo-BRD4-NUT “all” (purple), “inside” (blue), and “outside” (red) JF646 labeled single molecules, and TMR labeled whole condensates (orange) over a time range of 0.05 to 0.5 s (exposure time = 50 ms). For each sample, *n* = 30 cells. ***, *p* = 1.0 × 10^−15^ for “inside” versus “whole condensates”; *p* = 5.8 × 10^-13^ for “inside” versus “outside”; *p* = 8.5 × 10^-10^ for “inside” versus “all” single molecules by two-sided Kolmogorov-Smirnov test. Note that the MSD of “inside” molecules is significantly smaller than that of both “outside” and “all”, but larger than “whole condensates”.

**Fig. S6.**
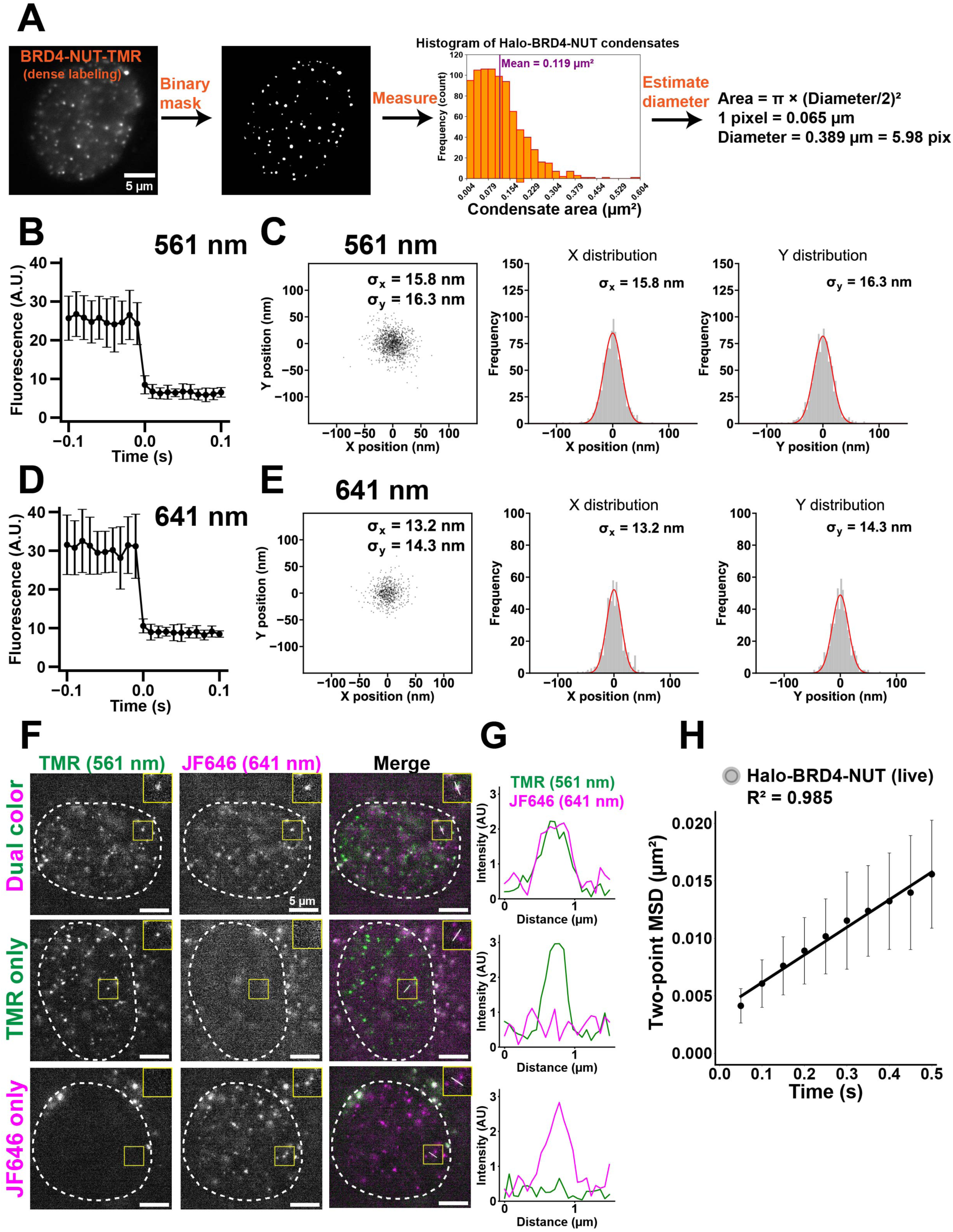
Single-molecule dual-color imaging of BRD4-NUT with a beam splitter system. **(A)** Workflow of calculating the dimensions of the Halo-BRD4-NUT condensates, as seen in live cells with dense TMR labeling. The image is binarized to obtain crisp outlines of the condensates. Then, the mean condensate area (purple line) is determined from the histogram of size distributions (orange bars). The condensate diameter is estimated by approximating them as circles: Area = π × (Diameter / 2)^2^. **(B)** Single-step photobleaching of ten representative Halo-TMR-BRD4-NUT single molecules imaged with a beam splitter system. The horizontal axis shows the time before and after photobleaching. **(C)** The position determination accuracy of Halo-TMR-BRD4-NUT single molecules imaged with a beam splitter system. Distribution of single-molecule displacements from the centroids in the (*x*, *y*)-plane in the 10-ms interval, *n* = 10 dots in FA-fixed HeLa cells. SD*_x_* and SD*_y_* of fitted Gaussian functions were 15.8 nm and 16.3 nm for TMR dots. **(D)** Single-step photobleaching of ten representative Halo-JF646-BRD4-NUT single molecules imaged with a beam splitter system. The horizontal axis shows the time before and after photobleaching. **(E)** The position determination accuracy of Halo-JF646-BRD4-NUT single molecules imaged with a beam splitter system. Distribution of single-molecule displacements from the centroids in the (*x*, *y*)-plane in the 10-ms interval, *n* = 10 dots in FA-fixed HeLa cells. SD*_x_* and SD*_y_* of fitted Gaussian functions were 13.2 nm and 14.3 nm for JF646 dots. **(F-G)** Example nuclei of HeLa cells expressing Halo-BRD4-NUT labeled with both TMR and JF646 (Dual color), TMR only, or JF646 only, as seen through the beam splitter. Representative single molecules are selected in yellow boxes and shown separately (top right corner). White punctate lines delineate the nuclei. The intensity line profiles for TMR and JF646 across the white lines in yellow boxes in (F) are plotted in (G). Note that our beam splitter system has no crosstalk between the TMR and JF646 signals. **(H)** Two-point MSD plot (mean ± SD among cells) between JF646- and TMR-labeled Halo-BRD4-NUT single molecules in HeLa cells from 0.05 to 0.5 s (tracked at 50 ms/frame, *n* = 29 cells).

**Fig. S7.**
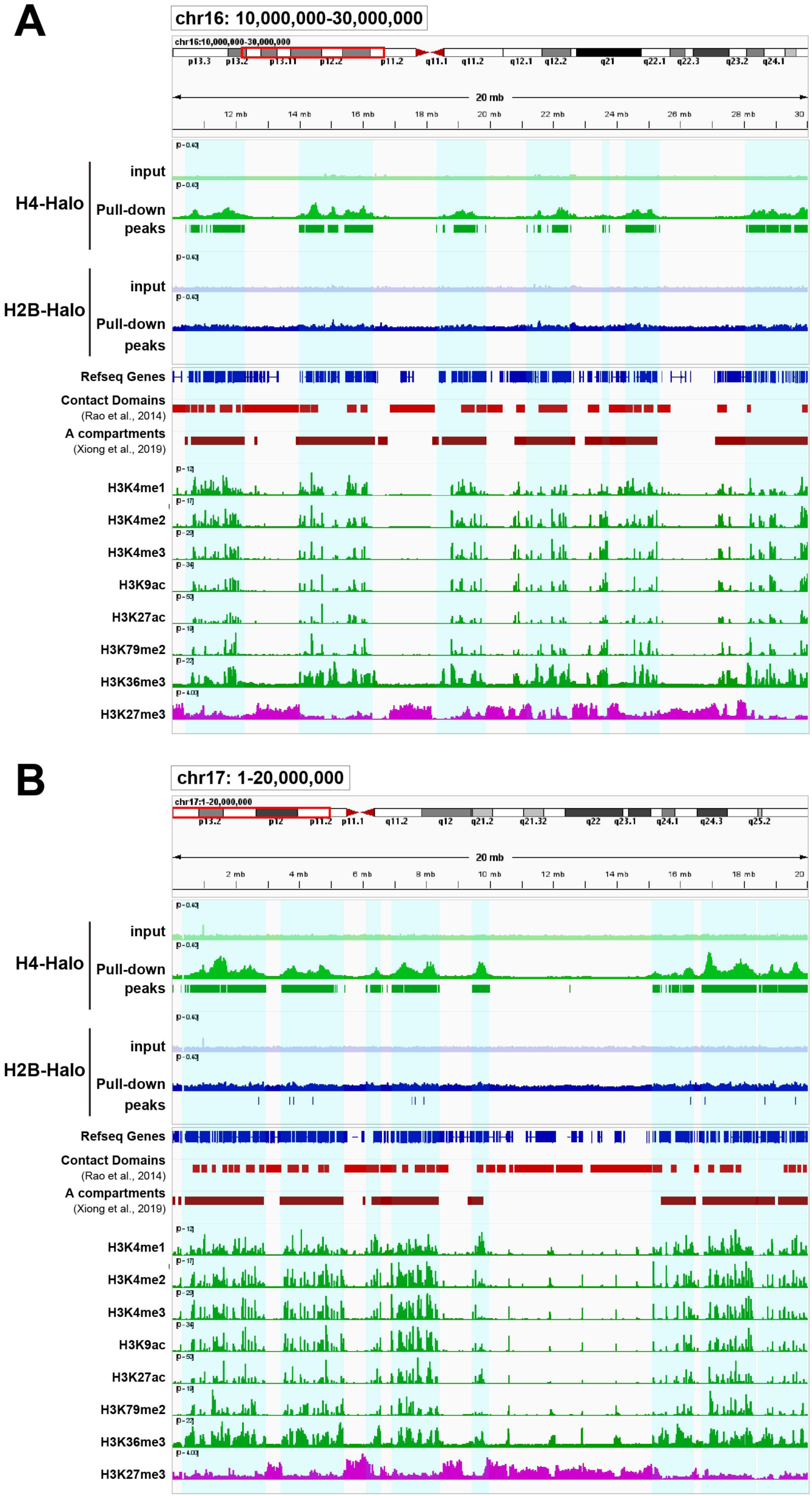
Visualization of H4-Halo labeled genomic regions. **(A and B)** Examples of H4-Halo– labeled regions on chromosome 16_10,000,000-30,000,000 (A) and chromosome 17_1-20,000,000 (B) using the Integrative Genomics Viewer (IGV) Browser. Blue shades on the background indicate the detected H4-HaloTag peak regions. The contact domains ^69^, predicted A-compartments ^70^, and active/repressive histone marks ^71^ are shown at the bottom for comparison.

**Fig. S8.**
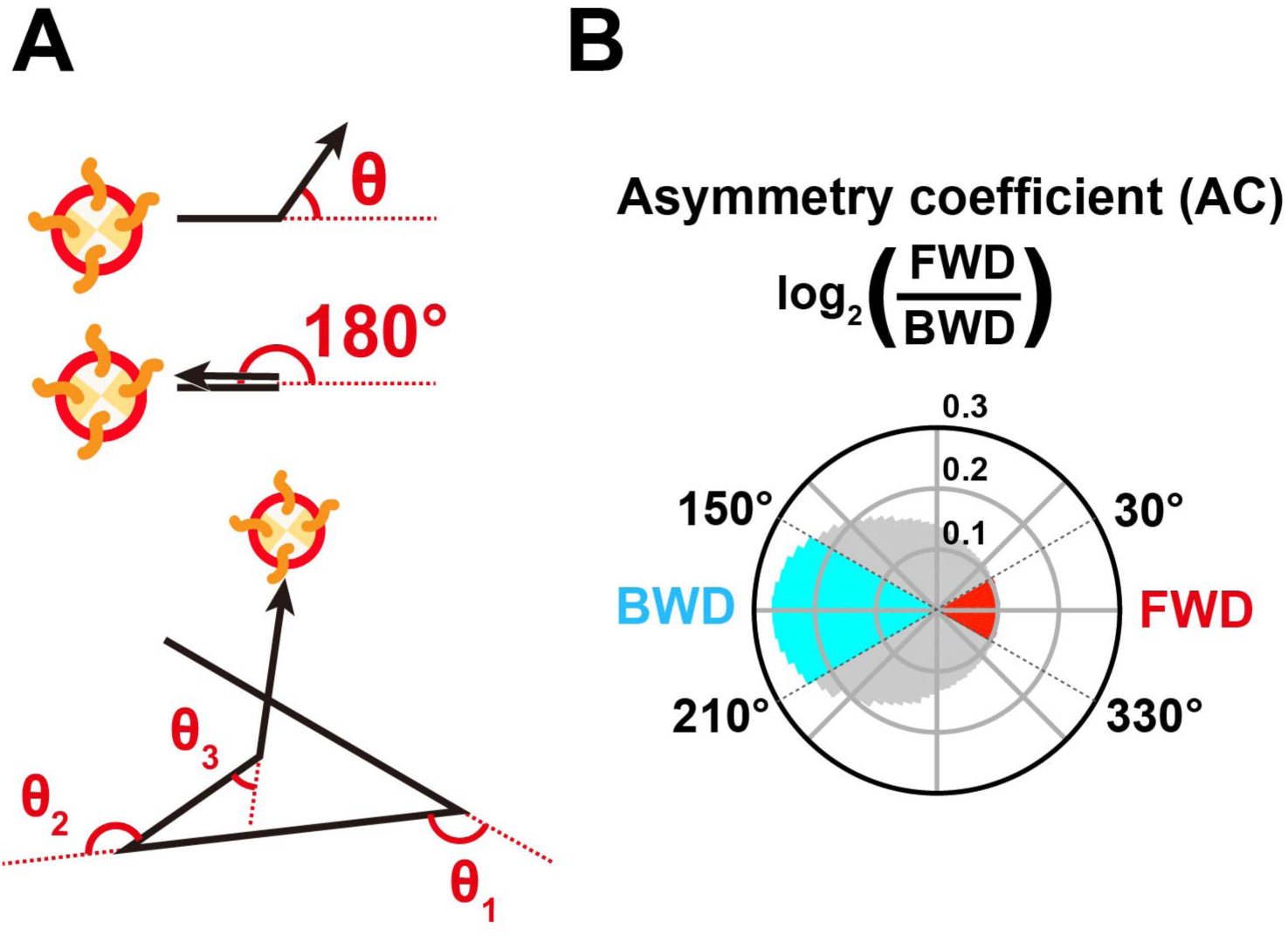
Nucleosome motion angle-distribution analysis in living cells. **(A)** Schematic for angle-distribution analysis. **(B)** Schematic for asymmetry coefficient (AC). AC was calculated as the logarithm to the base of 2 of the ratio between the frequencies of forward (FWD, −30° (330°) to +30°) and backward (BWD, 150° to 210°) angles. AC shows the deviation from a homogeneous distribution and is negative for angular distributions where the backward angles are dominant.

**Fig. S9.**
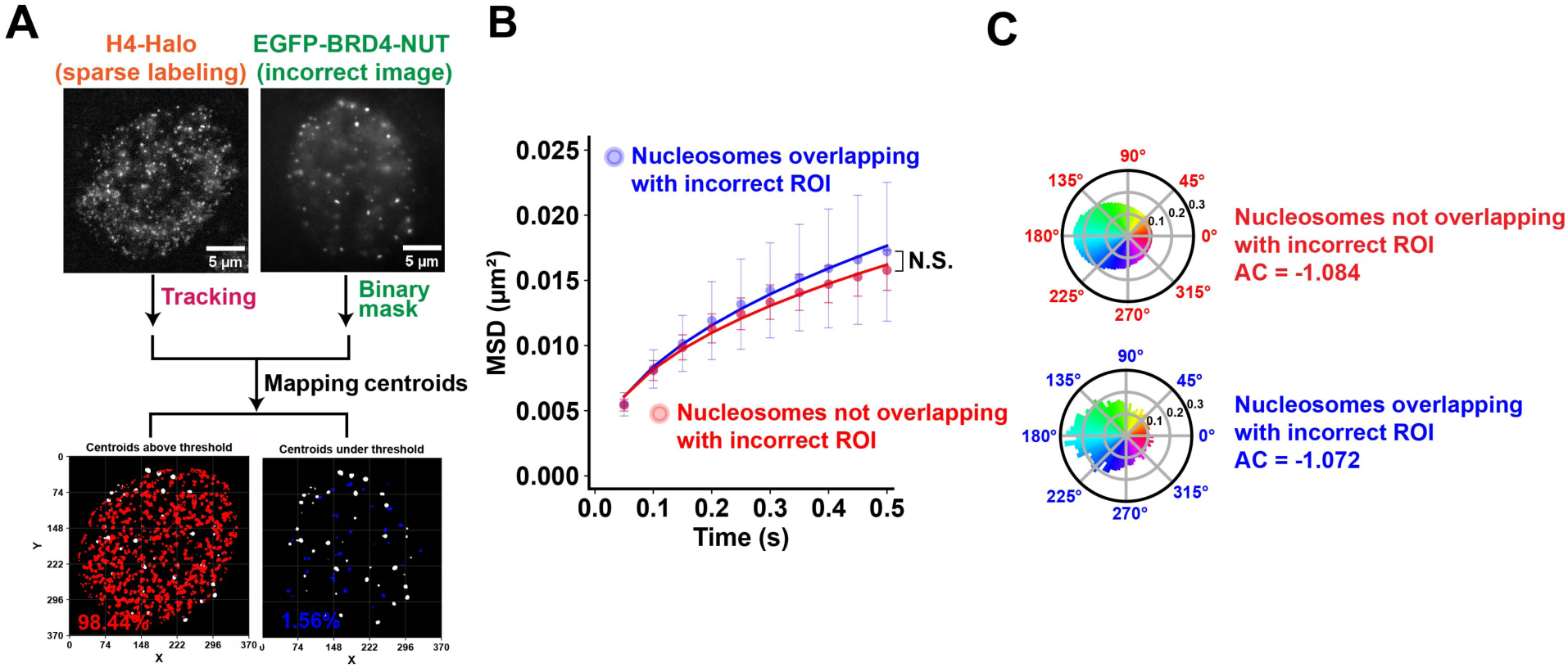
Validation of centroids mapping method using random binary masks. **(A)** Simplified workflow from Fig. 4C optimized for mapping H4-Halo trajectories on incorrect binary masks of EGFP-BRD4-NUT images. Top left: sparsely labeled H4-Halo image after background subtraction. Top right: EGFP-BRD4-NUT image in a different cell from top left. Bottom left: centroids of “not overlapping” H4-Halo trajectories overlayed on the incorrect binary mask. % - their percentage from all trajectories. Bottom right: centroids of “overlapping” H4-Halo trajectories overlayed on the incorrect binary mask. % - their percentage from all trajectories. **(B)** MSD plots (± SD among cells) of H4-Halo nucleosomes overlapping (blue) or not overlapping (red) with the incorrect masks. For each sample, *n* = 25 cells. N.S. (not significant), *p* = 0.156 for “overlapping” versus both “not overlapping” and MSD calculated from “all” H4-Halo trajectories by two-sided Kolmogorov-Smirnov test. **(C)** Measured angle distribution of “overlapping” and “not overlapping” nucleosomes (608,529 total number of angles for “not overlapping”; 14,158 angles for “overlapping”).

**Fig. S10.**
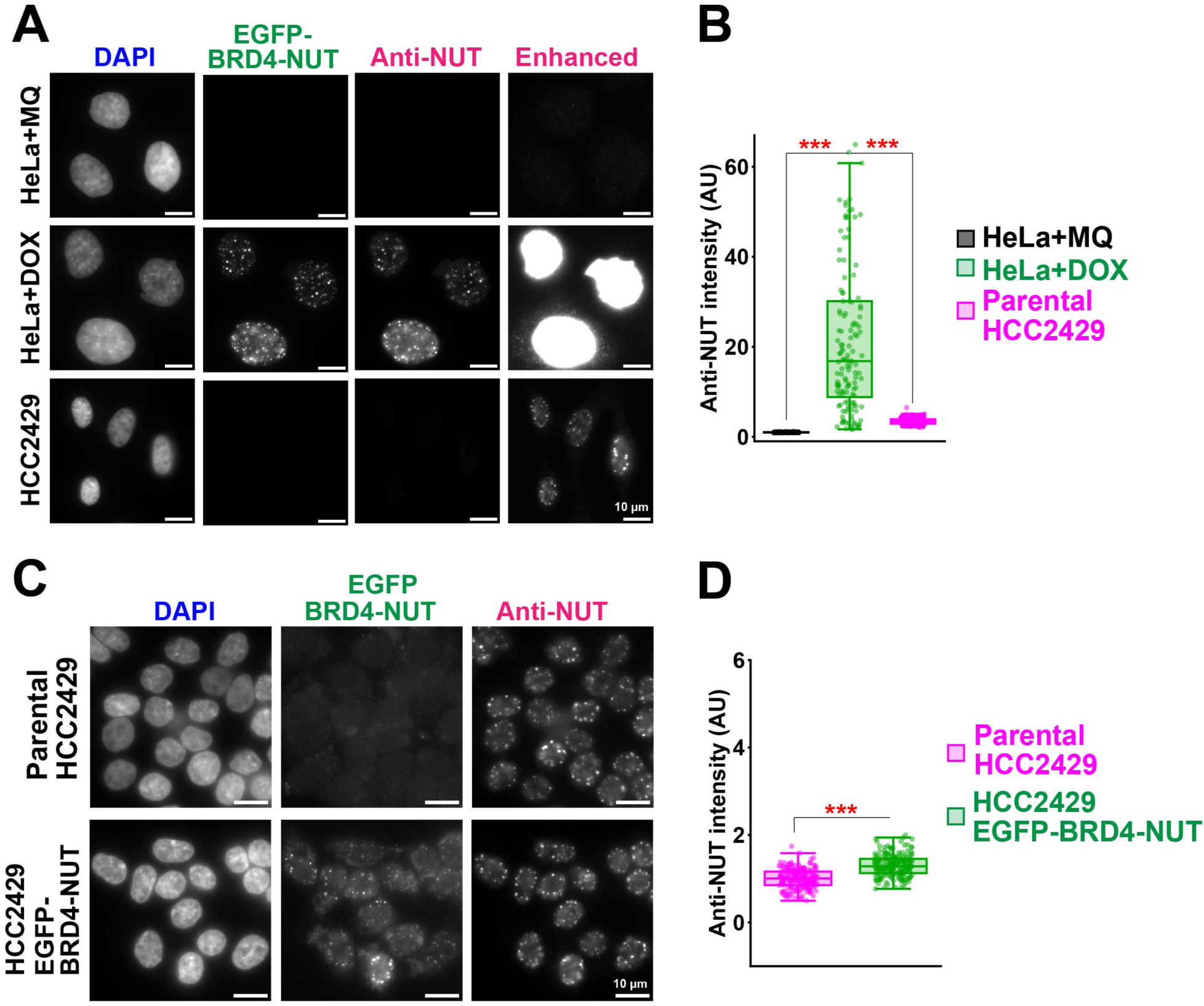
BRD4-NUT condensates are more strongly expressed in HeLa than in NC cells. **(A)** Immunostaining against NUT (Anti-NUT) in HeLa cells treated either with 0.1% Milli-Q (MQ) water or 1 µg/mL doxycycline (DOX) and HCC2429 NC cells. Note that the Anti-NUT foci are much weaker in HCC2429 cells, compared to HeLa + DOX, and are only observed when sufficiently enhanced (Enhanced). Scale bars, 10 μm. Image intensities are on the same scale for each column. **(B)** Boxplots of mean Anti-NUT signal intensities in HeLa + MQ, HeLa + DOX, and HCC2429 cells. Whiskers and bounds of boxes show maxima, minima, and first/third quartiles. The lines inside the boxes indicate the median values. Note a significant (∼6.7 times) difference in intensities between HeLa + DOX and HCC2429 cells. ***, *p* = 4.3 × 10^−47^ for HeLa + MQ versus HeLa + DOX; *p* = 1.1 × 10^−35^ for HeLa + DOX versus HCC2429 by two-sided Wilcoxon rank sum test. **(C)** Immunostaining against NUT (Anti-NUT) in parental and EGFP-BRD4-NUT-expressing HCC2429 cells. Note that the Anti-NUT foci are comparable in intensity between parental and EGFP-BRD4-NUT cells. Scale bars, 10 μm. Image intensities are on the same scale for each column. **(D)** Boxplots of mean Anti-NUT signal intensities in arbitrary units (AU) in parental and EGFP-BRD4-NUT-expressing HCC2429 cells. Note a small (∼1.3 times) difference in intensity between parental and EGFP-BRD4-NUT-expressing HCC2429 cells. ***, *p* = 3.2 × 10^−28^ for the parental versus EGFP-BRD4-NUT by two-sided Wilcoxon rank sum test. Whiskers and bounds of boxes show maxima, minima, and first/third quartiles. The lines inside the boxes indicate the median values.

**Fig. S11.**
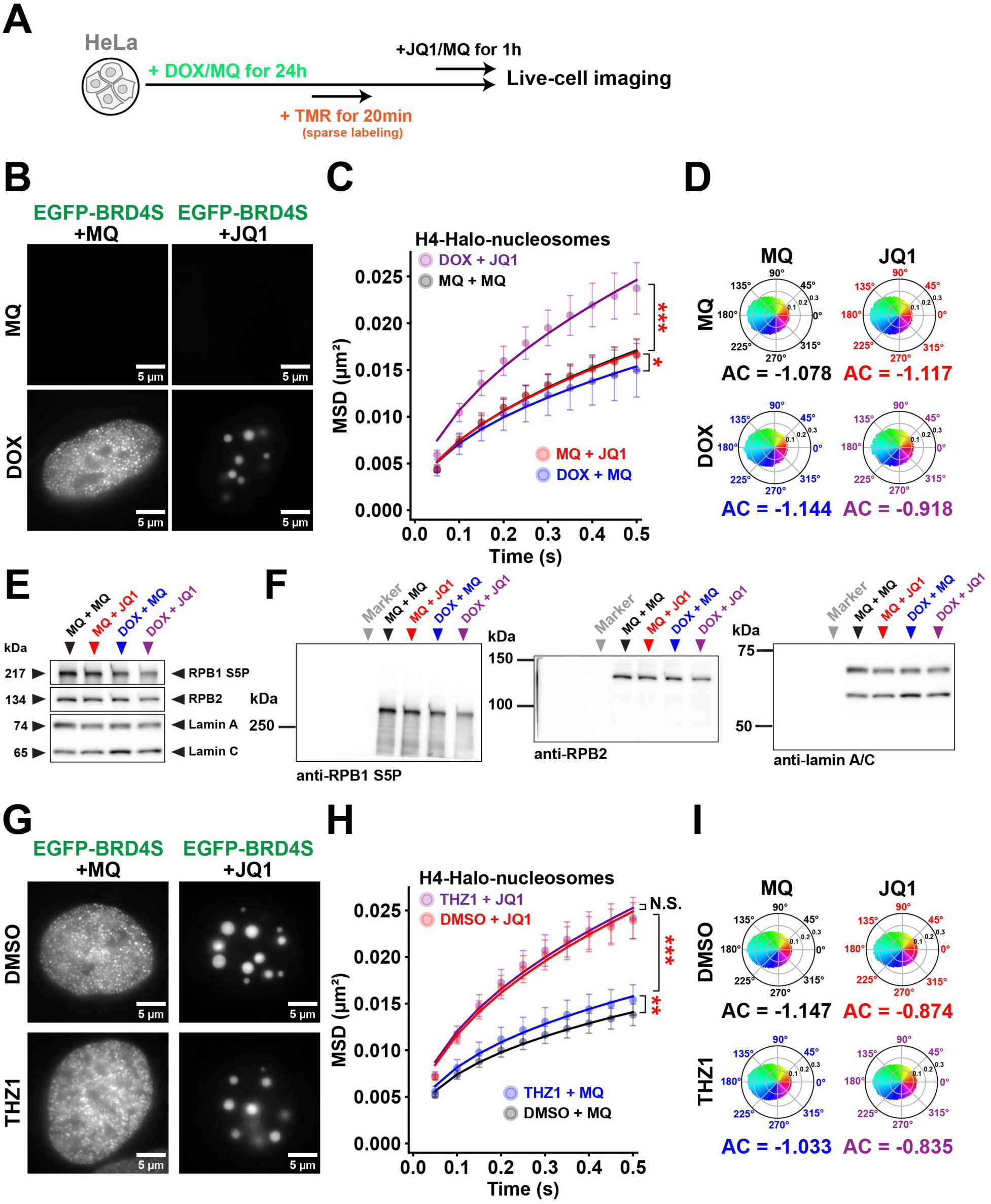
BRD4S overexpression and inhibition result in transcription inhibition. **(A)** Simplified protocol for 0.1% Milli-Q (MQ) water or 1 µM JQ1 treatment in the last 1 h and H4-Halo nucleosome sparse labeling during the 24-h incubation of cells in MQ or DOX. **(B)** Representative images of EGFP-BRD4S in cells treated with a combination of MQ/DOX and MQ/JQ1. Note that EGFP-BRD4S condensates are abundant, but small, in DOX + MQ. These coalesce into a few large spherical droplets in DOX + JQ1. Image intensities are on the same scale. **(C)** MSD plots (± SD among cells) of H4-Halo nucleosomes in MQ + MQ (black, *n* = 20 cells), MQ + JQ1 (red, *n* = 21 cells), DOX + MQ (blue, *n* = 19 cells), and DOX + JQ1 (purple, *n* = 20 cells) cells. ***, *p* = 3.0 × 10^−10^ for MQ + JQ1 versus DOX + JQ1; *, *p* = 4.9 × 10^−2^ for MQ + MQ versus DOX + MQ by two-sided Kolmogorov-Smirnov test. **(D)** Measured angle distribution of MQ + MQ, MQ + JQ1, DOX + MQ, and DOX + JQ1 nucleosomes (181,502 total number of angles for MQ + MQ; 173,938 angles for MQ + JQ1; 122,245 angles for DOX + MQ; 95,939 angles for DOX + JQ1). **(E)** Immunoblotting against RPB1 S5P, RPB2 (the second largest subunit of RNA Pol II), and lamin A/C in MQ + MQ, MQ + JQ1, DOX + MQ, and DOX + JQ1 HeLa cells. Note that the RPB1 S5P band is much weaker in lysates from DOX + JQ1 treated cells. **(F)** Uncropped images of the PVDF membranes used for immunoblotting in (E). Left: membrane incubated with anti-RPB1 S5P antibody, middle: anti-RPB2 antibody, or right: anti-lamin A/C antibody. **(G)** Representative images of EGFP-BRD4S overexpressed (+ DOX in all four) in cells treated with a combination of DMSO/THZ1 and MQ/JQ1. Note that EGFP-BRD4S condensates are abundant, but small, in both MQ conditions. These coalesce into a few large spherical droplets in both JQ1 conditions. In THZ1 + MQ, however, the condensates, while still small and abundant, appear more clustered. Image intensities were optimized for visualization. **(H)** MSD plots (± SD among cells) of H4-Halo nucleosomes in DMSO + MQ (black, *n* = 18 cells), DMSO + JQ1 (red, *n* = 21 cells), THZ1 + MQ (blue, *n* = 16 cells), and THZ1 + JQ1 (purple, *n* = 18 cells) cells. Note that EGFP-BRD4S was overexpressed (+ DOX) in all four cases. ***, *p* = 9.1 × 10^−10^ for THZ1 + JQ1 and THZ1 + MQ; **, *p* = 2.2 × 10^−3^ for THZ1 + MQ and DMSO + MQ; N.S. (not significant), *p* = 0.990 for DMSO + JQ1 versus THZ1 + JQ1 by two-sided Kolmogorov-Smirnov test. **(I)** Measured angle distribution of DMSO + MQ, DMSO + JQ1, THZ1 + MQ, and THZ1 + JQ1 nucleosomes (375,580 total number of angles for DMSO + MQ; 254,903 angles for DMSO + JQ1; 373,660 angles for THZ1 + MQ; 326,428 angles for THZ1 + JQ1). Note that EGFP-BRD4S was overexpressed (+ DOX) in all four cases.

## Movies S1–S8 legends

### Movie S1

Left: movie (10 ms/frame) of single molecules of Halo-JF646-BRD4-NUT in a living HeLa cell recorded by the sCMOS ORCA-Fusion BT camera (Hamamatsu Photonics). Note that clear and well-separated dots are visualized with single-step photobleaching profiles (Fig. 2D), suggesting that each dot represents a single Halo-JF646-BRD4-NUT molecule. Right: movie (10 ms/frame) of condensates of Halo-TMR-BRD4-NUT in the same cell. Note that the condensates also fluctuate in this time scale. Scale bar: 5 µm.

### Movie S2

Left: movie (50 ms/frame) of single molecules of Halo-JF646-BRD4-NUT in a living HeLa cell. Right: movie (50 ms/frame) of condensates of Halo-TMR-BRD4-NUT in the same cell. Scale bar: 5 µm.

### Movie S3

Movie (10 ms/frame) of the JF646- and TMR-labeled single molecules of Halo-BRD-NUT in a living HeLa cell. The two channels were recorded simultaneously using W-VIEW GEMINI (Hamamatsu Photonics). Left: JF646-labeled molecules. Middle: TMR-labeled molecules. Right: the merged movie of JF646-(magenta) and TMR-(green) labeled molecules. The yellow arrow indicates an example pair of neighboring JF646- and TMR-molecules. Scale bar: 5 µm.

### Movie S4

Movies (50 ms/frame) of the JF646- and TMR-labeled single molecules of Halo-BRD-NUT in a living HeLa cell. Left: JF646-labeled molecules. Middle: TMR-labeled molecules. Right: the merged movie of JF646-(magenta) and TMR-(green) labeled molecules. The yellow arrow indicates an example pair of neighboring JF646- and TMR-molecules. Scale bar: 5 µm.

### Movie S5

Left: movie (50 ms/frame) of single nucleosomes of H4-Halo-TMR in a living HeLa cell. Right: movie (50 ms/frame) of H4-Halo-TMR single nucleosomes in a living HCC2429 NC cell. Scale bar: 5 µm.

### Movie S6

Left: movie (50 ms/frame) of single nucleosomes of H4-Halo-TMR in a living HeLa cell with the doxycycline-inducible expression of EGFP-BRD4S treated with 0.1% Milli-Q water for 24 h. Right: movie (50 ms/frame) of nucleosomes in an analogous cell treated with 1 µg/mL doxycycline for 24 h. Scale bar: 5 µm.

### Movie S7

Left: movie (50 ms/frame) of single nucleosomes of H4-Halo-TMR in a living HeLa cell expressing EGFP-BRD4S condensates treated with 0.1% DMSO in the last 8 h of induction. Right: movie (50 ms/frame) of nucleosomes in an analogous cell treated with 500 nM TSA in the last 8 h of induction. Scale bar: 5 µm.

### Movie S8

Left: movie (50 ms/frame) of single nucleosomes of H4-Halo-TMR in a living HeLa cell expressing EGFP-BRD4S condensates treated with 0.1% Milli-Q water in the last 1 h of induction. Right: movie (50 ms/frame) of nucleosomes in an analogous cell treated with 1 µM JQ1 in the last 1 h of induction. Scale bar: 5 µm.

